# The Unique Pt(II)-Induced Nucleolar Stress Response and its Deviation from DNA Damage Response Pathways

**DOI:** 10.1101/2024.06.05.597606

**Authors:** Hannah C. Pigg, Katelyn R. Alley, Christopher R. Griffin, Caleb H. Moon, Sarah J. Kraske, Victoria J. DeRose

## Abstract

The mechanisms of action for the platinum compounds cisplatin and oxaliplatin have yet to be fully elucidated, despite the worldwide use of these drugs. Recent studies suggest that the two compounds may be working through different mechanisms, with cisplatin inducing cell death via the DNA damage response (DDR) and oxaliplatin utilizing a nucleolar stress-based cell death pathway. While cisplatin- induced DDR has been subject to much research, the mechanisms for oxaliplatin’s influence on the nucleolus are not well understood. Prior work has outlined structural parameters for Pt(II) derivatives capable of nucleolar stress induction. In this work, we gain insight into the nucleolar stress response induced by these Pt(II) derivatives by investigating potential correlations between this unique pathway and DDR. Key findings from this study indicate that Pt(II)-induced nucleolar stress occurs when DDR is inhibited and works independently of the ATM/ATR-dependent DDR pathway. We also determine that Pt(II)-induced stress may be linked to the G1 cell cycle phase, as cisplatin can induce nucleolar stress when cell cycle inhibition occurs at the G1/S checkpoint. Finally, we compare Pt(II)-induced nucleolar stress with other small-molecule nucleolar stress-inducing compounds Actinomycin D, BMH-21, and CX-5461, and find that only Pt(II) compounds cause irreversible nucleolar stress. Taken together, these findings contribute to a better understanding of Pt(II)-induced nucleolar stress, its deviation from ATM/ATR-dependent DDR, and the possible influence of cell cycle on the ability of Pt(II) compounds to cause nucleolar stress.

## Introduction

Small-molecule platinum compounds are of great interest for their versatile ligand platforms and ability to undergo ligand exchange with biomolecules including RNA, DNA and proteins as well as metabolites. These properties underlie platinum-based chemotherapeutics, including cisplatin, carboplatin and oxaliplatin, which have been pivotal in the treatment of various types of cancers for nearly 50 years. Although these compounds have been widely utilized for frontline treatments, their clinical applications are often limited by severe drug side effects and the emergence of drug resistant cancers. Each of these three platinum drugs have been shown to have distinct anti-tumor activity profiles. For example, many colorectal cancer lines are resistant to treatment with cisplatin and carboplatin, while being sensitive to treatment with oxaliplatin. The mechanisms behind the variation in anti-tumor activity between platinum compounds is not well understood, and elucidating the mechanisms of action of these compounds is crucial for basic understanding of small molecule interactions as well as to the development of clinical treatments that can target drug resistant cancer lines (1–3). Until recently, it was thought that all three Pt(II) compounds caused cell death through the DNA damage response (DDR). However, over the last decade, new research has emerged indicating that while cisplatin and carboplatin do appear to elicit cell death through DDR, oxaliplatin may instead be inducing cell death through a transcription-translation inhibition-based pathway (4–6). In this pathway, ribosome biogenesis, which occurs in the nucleolus, is inhibited and cells display a nucleolar stress response, leading to cell death or senescence. The Pt(II)-induced nucleolar stress response is still not well characterized, and is an area of great interest for nucleolar targeting by small molecules as well as a basic understanding of nucleolar function (7, 8). Additionally, determining how Pt(II)-induced nucleolar stress functions in tandem to or independently of Pt(II)-induced DDR is of high importance to a comprehensive understanding of the mechanism of action of platinum compounds.

DDR is a complex cellular signaling pathway, which detects various forms of DNA damage and attempts to repair DNA prior to DNA synthesis and cell division. DDR is primarily initiated by sensor protein kinases: ataxia telangiectasia mutated (ATM) and ataxia telangiectasia and Rad-3 related (ATR). The type of DNA damage occurring dictates what sensor kinase is activated, where generally ATM is activated by DNA double-stranded breaks (DSBs) and ATR is activated by a variety of DNA damage inducing stimuli, including DSBs, DNA single-stranded breaks (SSBs) and DNA crosslinks (9). Activation of ATM/ATR will then trigger a cascade of signal transduction events which can be broadly classified into 5 primary DNA repair pathways: homologous recombination (HR) repair, nucleotide excision repair (NER), non-homologous end joining (NHEJ) repair, base excision repair (BER), and mismatch repair (MMR). The type of DNA damage dictates the repair pathway or pathways that are activated and will ultimately result in DNA repair, cell cycle arrest, or cell death via apoptosis (10, 11). The proper functioning of these DNA repair pathways is crucial for maintaining a stable genome and pathway defects have been shown to increase a person’s risk of developing various types of cancer (12–14). Given the significant role of DDR in genome stability and cell cycle regulation, many cancer treatments target DDR in order to propagate cell cycle arrest and cell death in cancer cells (15, 16).

Upon uptake, platinum compounds are known to enter the nucleus of cells and form intra- and inter-strand DNA adducts, triggering DDR. DDR activation in response to platinum drug treatment has been extensively studied, however the specific mechanisms and repair pathways involved in platinum-induced DDR have not been fully elucidated and vary based on the platinum compound and cell line (17–19). Therefore, understanding how the differing drug profiles of oxaliplatin and cisplatin may be linked to their formation of DNA-Pt adducts is of high importance (20–23). Both cisplatin and oxaliplatin have been shown to form DNA-Pt adducts, however cisplatin shows notably higher levels of DNA-Pt adduct formation across various cancer cell lines. Additionally, both platinum compounds have been shown to form adducts at the same sites on DNA, the N7 position of adjacent guanine residues, and show a preference of nuclear DNA over mitochondrial DNA (24, 25). Evidence suggests that DNA adduct accumulation does not likely reflect drug toxicity, as many cells show higher toxicity to oxaliplatin, despite having quantitatively lower levels of oxaliplatin-DNA adducts compared to cisplatin adducts (26). Therefore, the differences in drug cytotoxicity may instead be linked to differences in biological properties of the DNA adducts formed, where the bulky diaminocyclohexane (DACH) ligand present in oxaliplatin and not cisplatin may have significant influences (25).

The ability of cells to repair DNA damage induced by platinum compounds is an important factor in determining cytotoxicity, and differences in repair mechanism for cisplatin vs oxaliplatin DNA-adducts may in part help to explain there varying sensitivity across cancer cell lines (27). Various DNA repair pathways have been shown to be involved in repairing DNA lesions induced by platinum compounds, including NER, MMR, HR and NHEJ (17, 28). Studies have found that oxaliplatin and cisplatin elicit different expression profiles for genes associated with these DNA repair pathways across different cell lines (4, 29). Surprisingly, repair efficiency for oxaliplatin vs cisplatin DNA-adducts is generally not significantly different across studies, indicating that it is not likely the ability of DNA repair proteins to recognize oxaliplatin-DNA adducts that is eliciting its drastic differences in cytotoxicity compared to DNA-adduct concentration (30, 31). Given the various dissimilarities between oxaliplatin and cisplatin regarding genomic binding activity, cancers cell types they are most effective in treating, and physiological side effects induced by each drug, it is not surprising that recent work has focused on the potential of oxaliplatin working through a different cell death mechanism to that of cisplatin.

Recent studies focused on gaining a better understanding of oxaliplatin’s mechanism have been primarily based on the compounds’ ability to cause ribosome biogenesis inhibition, rather than or in addition to DDR (4, 8). The ability of oxaliplatin to effectively cause inhibition of ribosome biogenesis at clinically relevant concentrations was determined over 15 years ago, when it was demonstrated by Burger, et. al to inhibit the production of the 47 S rRNA precursor at concentrations as low as 3 µM in the human sarcoma cell line 2FTGH. In this study, cisplatin was also tested, but had to be used at 50 µM concentrations or higher to induce the same levels of inhibition as oxaliplatin (32). Drugs that inhibit ribosome biogenesis have been used to treat cancer since as early as 1954, with the use of the anticancer drug Actinomycin D (ActD) (33). Ribosome biogenesis is the process by which ribosomal DNA (rDNA) is transcribed into ribosomal RNA (rRNA) by RNA Polymerase I (Pol I) and then further processed into ribosomal subunits, which eventually become mature ribosomes (34). This process primarily takes place in the cellular nucleolus, a membraneless sub-section of the nucleus. The nucleolus has emerged as an organelle harboring much interest from researchers in the last decade, as it has been shown to not only be involved in ribosome biogenesis, but also to have multiple other cellular functions in regulating cell cycle and cellular homeostasis (35, 36). Any disruption to the integrity or function of the nucleolus can be referred to as nucleolar stress.

Previous work from our lab has focused on gaining a better understanding of nucleolar stress induced by oxaliplatin and other Pt(II) compounds that were found to induce nucleolar stress (**Figure 1**). Our work found that nucleolar stress induced by platinum compounds is highly specific and is sensitive to small changes in platinum compound structure and aromaticity. This stress response is also time-dependent and is accompanied by rRNA transcription inhibition as early as 3 hours after drug treatment. Additionally, there is no correlation found between Pt(II)-induced nucleolar stress and whole-cell or nucleolar drug uptake, or platinum DNA adduct formation (5, 37–41). Recently, it has been suggested that Pt(II)-induced nucleolar stress may be initiated by ATM/ATR-dependent DDR. Specifically, it was shown that when ATM or ATR were inhibited, rRNA transcription inhibition by oxaliplatin decreased, indicating the potential role of these proteins in the nucleolar stress response pathway. However, these results show that ATM/ATR inhibition did not fully combat rRNA transcription inhibition by oxaliplatin, as significant inhibition was still observed in the presence of inhibitors, suggesting that ATM/ATR-dependent DDR is not solely responsible for the nucleolar stress response (42). Other work suggests that Pt(II)-induced nucleolar stress may be a product of liquid-liquid demixing of the nucleoli upon oxaliplatin treatment, leading to rRNA transcription shutdown and eventual cell death (43). Overall, the process by which oxaliplatin and other nucleolar stress inducing derivatives induce stress is still not well understood and warrants further studies into the mechanism.

**Figure 1:**
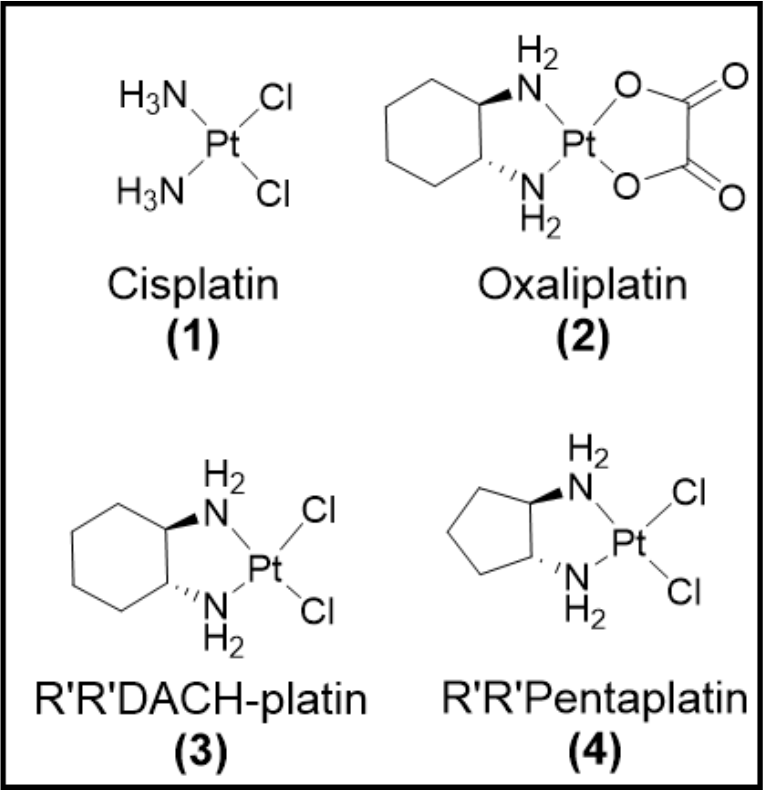
Pt(II) compounds utilized in this study. Cisplatin (CisPt) 1 does not induce nucleolar stress, whereas Oxaliplatin (OxaliPt) 2 and derivatives R’R’DACH-platin (DACH-Pt) 3, and R’R’Pentaplatin (Penta-Pt) 4 induce nucleolar stress.

In this manuscript, we further investigate the potential roles that DDR, cell cycle arrest, cell type, and compound treatment time may have on the unique Pt(II)-induced nucleolar stress pathway. Additionally, we study the reversibility of Pt(II)-induced nucleolar stress and compare this to other small-molecule nucleolar stress inducing compounds to help further elucidate the potential mechanisms involved in the pathway. Our findings suggest that although oxaliplatin and derivatives can initiate the DDR pathway, there does not appear to be a significant link between ATM/ATR-dependent DDR and Pt(II)-induced nucleolar stress, and nucleolar stress can occur independent of ATM/ATR-dependent DDR activation. Additionally, we determine a potential link between G1 cell cycle arrest and Pt(II)-induced stress, as cisplatin is found to cause nucleolar stress when the G1/S checkpoint is inhibited. We also conclude that unlike other small-molecule nucleolar stress-inducing compounds, Pt(II)-induced stress is non-reversible upon compound removal, suggesting it is not working through the same mechanisms as these organic compounds. Finally, many of these findings hold true across various treatment times and cell lines, differing in tumor type, drug sensitivity profiles, gene expression patterns, and cell cycle progression times (44–46). Taken together, these findings contribute to a better understanding of Pt(II)-induced nucleolar stress and its deviation from ATM/ATR-dependent DDR, and further elucidates how oxaliplatin’s unique ribosome biogenesis inhibition pathway differs from the DDR dependent pathway induced by cisplatin (4, 43, 47).

## Results/Discussion

### Oxaliplatin and derivatives do not cause significant activation of key DDR initiation proteins

In order to investigate differences in DDR activation between cisplatin and oxaliplatin, as well as other nucleolar stress-inducing Pt(II) derivatives (**Figure 1**), we investigated proteins involved in the initiation of Pt(II)-induced DDR. Although various pathways have been shown to be involved in platinum-induced DNA damage repair, BER, NER, and HR are often credited as the primary repair pathways for Pt-adducts (48–50). The initial cellular response to DNA damage is controlled by the phosphoinositide-3-kinase-related protein kinase (PIKK) family, which includes the Ataxia telangiectasia- and Rad3- related (ATR) and Ataxia telangiectasia mutated (ATM) kinases. Several studies support the role of ATR and ATM in the initiation and regulation of BER, NER, and HR (51–53). Additionally, the histone H2AX is a substrate for both ATM and ATR and has been shown to activate with the initiation of BER, NER and HR in response to oxaliplatin and cisplatin treatments (54–56). Given the overlapping role of ATM, ATR, and H2AX among the three pathways, we chose to focus primarily on these proteins for our studies focused on DDR and Pt(II)-induced nucleolar stress.

### Stress-inducing Pt(II) compounds elicit lower H2AX activation levels compared to cisplatin

H2AX phosphorylation is frequently utilized as a tool to determine the ability of treatments to induce DDR and the capacity of cell lines to repair damaged DNA (57, 58). Previous work has shown that at short drug treatment times, CisPt causes higher activation of H2AX compared to OxaliPt across cell lines (4, 38, 42). Here we extend these studies by examining additional cell lines and other stress-inducing Pt(II) compounds, DACH-Pt and Penta-Pt, as well as looking at longer drug treatment times. DACH-Pt is used to eliminate the slower ligand exchange kinetics observed with OxaliPt (59), and the smaller ring size of Penta-Pt gives it an intermediate effect on nucleolar stress (5, 37). A549 cells have been shown to undergo nucleolar stress after treatment with OxaliPt, DACH-Pt, and Penta-Pt, but show minimal stress induction from CisPt treatment (5, 37). In addition to A549 cells, U-2 OS and HCT116 cells were chosen as they are well characterized in the literature and show differences in their drug sensitivity profiles and gene expression patterns (43, 45, 60, 61). Specifically, HCT116 cells often show higher sensitivity to oxaliplatin and lower sensitivity to cisplatin, whereas U-2 OS and A549 cells show lower sensitivity to oxaliplatin and higher sensitivity to cisplatin. Treatment times of 3 and 5 hours were chosen in order to investigate DDR activity during times when Pt(II)-induced nucleolar stress is first observed for OxaliPt and DACH-Pt treatments, and 24 hour treatments were used to monitor trends in activation over time.

Figure 2 shows a comparison of H2AX activation levels at various drug treatment times in A549 cells (Figure 2A), U-2 OS cells (Figure 2B), and HCT116 cells (Figure 2C). At both short and extended drug treatment times, OxaliPt treatment results in significantly lower levels of H2AX activation in comparison with CisPt. In A549 cells (Figure 2A), DACH-Pt and Penta-Pt treatment induces higher degrees of H2AX activation compared to OxaliPt, and more closely resemble the activation profile for CisPt at the 3- and 5-hour drug treatment time points. This initial increase in H2AX activation may be attributed to the labile chloride ligands of DACH-Pt and Penta-Pt, which afford faster aquation rates compared to the oxalate ligand of OxaliPt. Interestingly, by 24 hours of drug treatment, H2AX activation levels from both DACH-Pt and Penta-Pt decrease to levels comparable to those of OxaliPt, while CisPt shows a significant increase in activation compared to shorter drug treatment times. These results suggest that the DNA damage induced by DACH- Pt and Penta-Pt at short time points can be repaired by 24 hours, while DNA damage from CisPt at 24 hours is still significant and DNA repair is minimal. OxaliPt activation levels stay relatively the same across all 3 drug treatment times in A549 cells.

**Figure 2:**
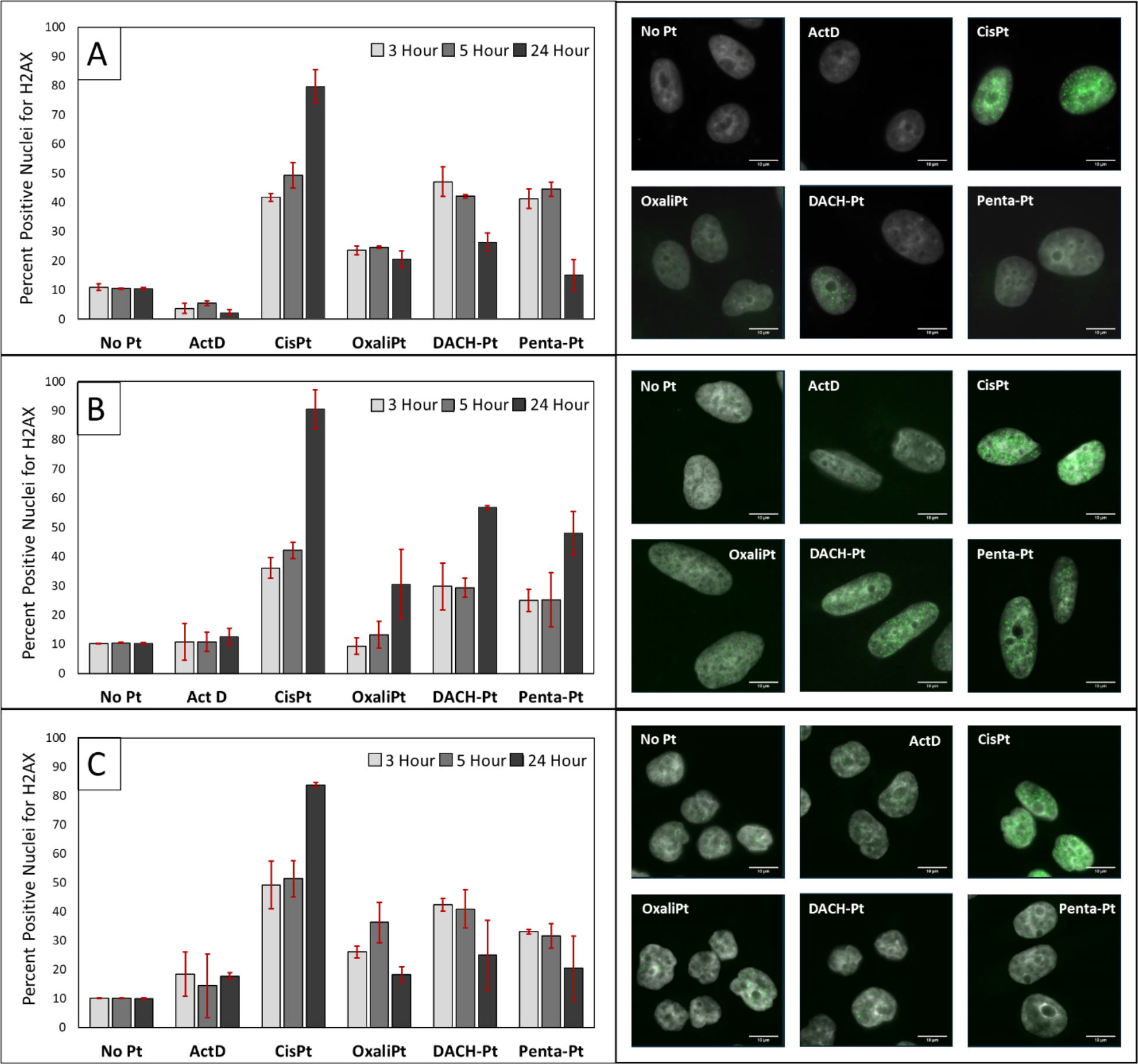
Q**u**antification **of H2AX activation after 3, 5 and 24 hr. drug treatments.** Cells were treated with 10 µM platinum compounds or 5 nM of ActD for 24 hours in A549 (**A**), U-2 OS (**B**), and HCT116 (**C**) cell lines. The immunofluorescence intensity for ƴH2AX was then measured and percent positive nuclei were determined, where a positive threshold was determined by the 90th percentile of the untreated control (see Methods). The data points represent the average percent positive nuclei across three separate biological replicates with standard deviation. Representative cell images of the 24 hr. treatment are shown with ƴH2AX (green) overlayed with DAPI (grey) for each cell line.

Compared to A549 cells, U-2 OS cells (Figure 2B) had lower activation levels for all Pt(II) compounds after 3- and 5-hour drug treatments, but still showed similar trends between compounds at these time points. Unlike A549 cells, at 24 hours of drug treatment, there was an increase in H2AX activation for all the nucleolar stress-inducing compounds compared to earlier time points. Interestingly, the percent H2AX activation at 24 hours for the stress-inducing compounds in U-2 OS cells were similar to the levels observed for the 5-hour drug treatments in A549 cells. Given this observation, we questioned if the Pt(II) compounds were initiating a slower response in U-2 OS cells compared to A549 cells and if increasing the drug treatment time would lead to a decrease in activation by nucleolar stress-inducing compounds. At 48-hour drug treatment times (**Figure S1**) U-2 OS cells show trends similar to the 24-hour treatments in A549 cells, with OxaliPt, DACH-Pt and Penta-Pt decreasing in activation and CisPt showing higher levels of H2AX activation. These results suggest that H2AX activation in U-2 OS cells may occur more slowly than in A549 cells, but that DNA damage induced by stress causing compounds is still able to be repaired, compared to DNA damage induced by CisPt. This may be caused by the slower doubling time for U-2 OS cells of ∼36 hours, compared to A549 and HCT116 cells which have a doubling time of ∼18 hours (62). Cells with slower proliferation rates spend less time in the S-phase, where DDR activation primarily occurs, and previous studies have shown that proliferation rate may have some effect on cellular responses to cisplatin and other chemotherapeutics (63, 64).

HCT116 cells (Figure 2C) show similar results to A549 cells, having higher H2AX activation by DACH-Pt and Penta-Pt at short time points and a decrease in activation for these compounds at 24-hour drug treatments. Likewise, by 24-hour drug treatment, CisPt shows higher levels of activation.

Taken together, these results suggest that nucleolar stress-inducing Pt(II) compounds cause overall lower activation of H2AX compared to CisPt across multiple cell lines. Additionally, although there is an initial increase in H2AX activation for some of the nucleolar stress-inducing compounds, by 24 hours the DNA damage caused by these compounds appears to be more easily repaired than damage caused by CisPt.

### DNA damage by nucleolar stress-inducing Pt(II) compounds leading to H2AX activation may be more easily repaired in comparison with cisplatin

To further explore DNA damage repair between cisplatin and stress inducing compounds, we measured H2AX activation after an initial drug treatment, followed by a drug free recovery period (Figure 3). We hypothesized that if DNA damage by nucleolar stress causing compounds was more easily repaired, we would see a larger decrease in activation for these compounds after the drug was removed in comparison with. For these studies A549 cells were treated with compounds for 5 hours followed by a chase with fresh drug-free media for 24 hrs. Following the recovery period, H2AX levels were quantified and then compared to levels observed following 5 hours of drug treatment with no drug free chase period. H2AX activation levels significantly decreased for nucleolar stress-inducing compounds after the 24-hour drug free recovery period, while CisPt treatment induced even higher levels of activation after the recovery period. This further indicates that H2AX activating DNA damage induced by nucleolar stress-inducing compounds may be repaired more quickly than CisPt-induced DNA damage. We next wanted to determine if the trends seen for H2AX extended to other key DDR initiation proteins.

**Figure 3:**
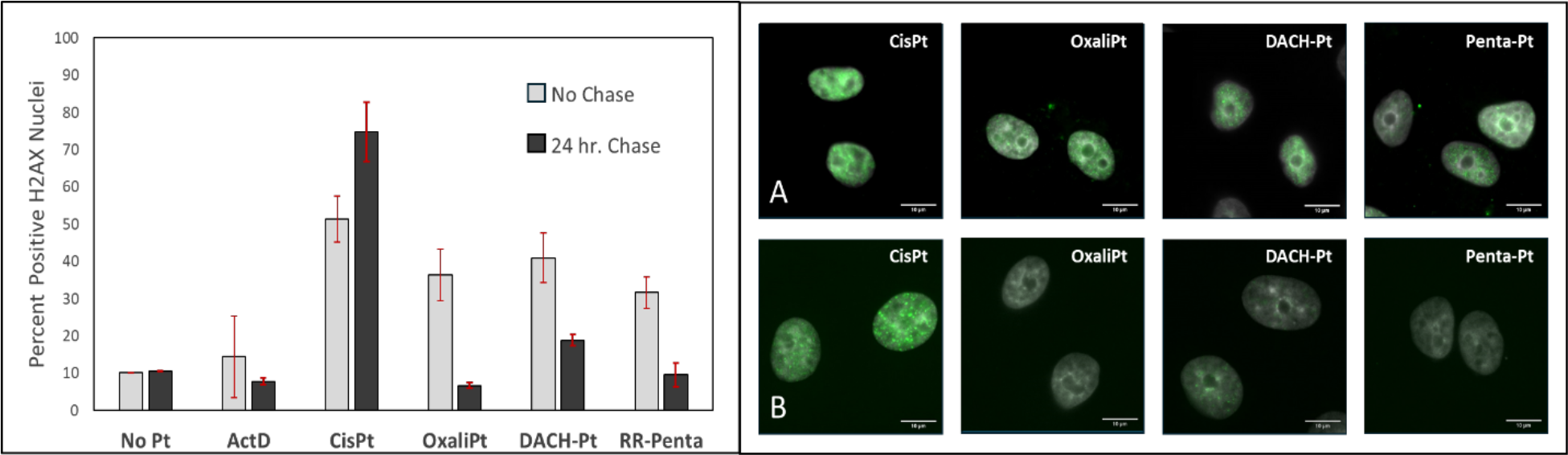
Q**u**antification **of H2AX activation after 5 hr. drug treatment and a 24 hr. drug free chase period.** Cells were treated with 10 µM platinum compounds or 5 nM of ActD for 5 hours in A549 cells. Following treatment, drug-free media was replaced on cells for 24 hours. The immunofluorescence intensity for ƴH2AX was then measured and quantified using the same method described in **Figure 2**. The data points represent the average percent positive nuclei with standard deviations across three separate biological replicates. Representative cell images of ƴH2AX (green) overlayed with DAPI (grey) are shown for each cell line after 5 hr. drug treatment alone (**A**), and 5 hr. drug treatment followed by 24 hr. drug free chase (**B**).

### Nucleolar stress-inducing Pt(II) compounds cause lower activation of ATM compared to cisplatin and other DDR inducing compounds

In addition to H2AX, the kinases proteins ATM and ATR are also critical in the initial detection of DNA damage and the initiation of the DDR. To determine if the distinct differences observed for nucleolar stress-inducing Pt(II) compounds in our H2AX studies extended to other key DDR initiation proteins we next aimed to measure the activation levels of ATM and ATR after drug treatment. We initially hoped to monitor ATR phosphorylation levels but found that all three cell lines had constitutively activated ATR and we observed no measurable increase of ATR phosphorylation with any platinum compound treatment or the control DNA damage reagent bleomycin (data not shown). Thus, we did not include ATR activation in these initial studies. Compared to ATM, ATR can be activated by a larger array of stimuli, including replicative stressors, and is frequently activated in cancer cell lines, as they often exhibit high degrees of replicative stress (65, 66). HCT116 cells were also omitted from these studies, as ATM activation was not observed for any of the given drug treatments nor for the positive control treatment with bleomycin, a DDR inducing compound known to cause ATM phosphorylation (67). The lack of activation in HCT116 cells may be a result of these cells having downregulated levels of the ATM protein, which has previously been observed across various colorectal cancer cell lines (68, 69). In contrast, A549 and U-2 OS cells treated with bleomycin showed significant ATM activation after only 3 hours of drug treatment (Figure 4).

**Figure 4:**
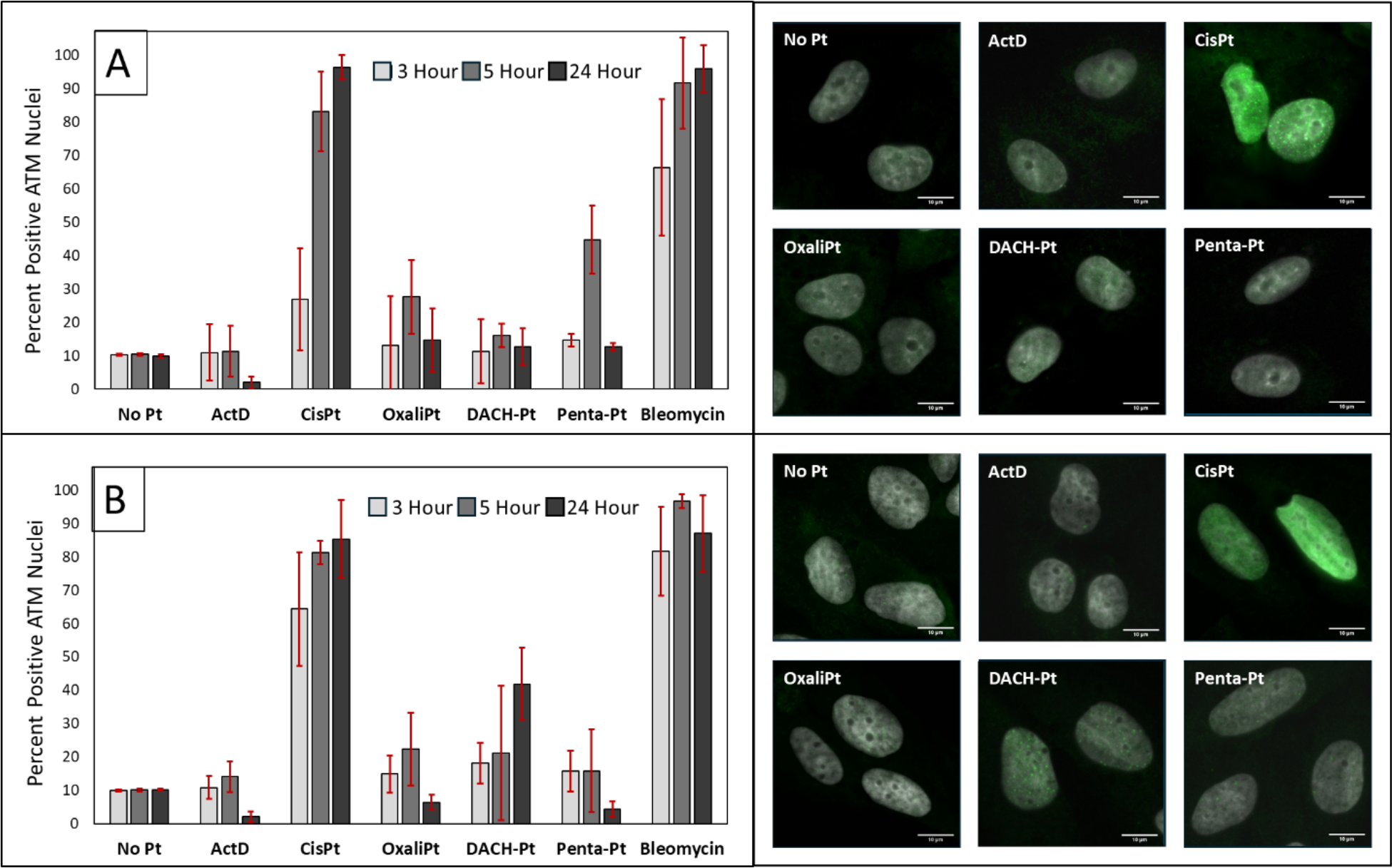
Q**u**antification **of ATM activation after 3, 5 and 24 hr. drug treatments.** Cells were treated with 10 µM platinum compounds or 5 nM of ActD for 24 hours in A549 (**A**) and U-2 OS (**B**) cell lines. The immunofluorescence intensity for pATM was then measured and percent positive nuclei were determined, where a positive threshold was determined by the 90th percentile of the untreated control. The data points represent the average percent positive nuclei with standard deviations across three separate biological replicates. Representative cell images for each cell line after 24 hr. drug treatment are shown with pATM (green) overlayed with DAPI (grey).

We measured the phosphorylation of ATM (pATM) in A549 and U-2 OS cells after 3, 5 and 24 hours of drug treatment (Figure 4). In A549 cells (Figure 4A) and U-2 OS cells (Figure 4B), ATM activation levels mimicked trends observed for H2AX activation. Treatments with CisPt, and DDR inducer bleomycin, resulted in an increase in ATM activation levels as drug treatment time increased. As with H2AX activation, 24-hr drug-free chase did not reduce ATM activation following cisplatin treatment (**Figure S2**). By contrast, the nucleolar stress inducing compounds only induce ATM activation levels slightly higher than that of the no drug controls (Figure 4), indicating that DNA damage leading to activation of ATM may not be occurring at significant levels. Results from these studies further support the claim that nucleolar stress inducing compounds appear to have an ATM phosphorylation profile distinct from known DDR inducing compounds.

### Pt(II)-induced nucleolar stress can occur in the absence of DDR activation

#### Nucleolar Stress Inducing Compounds Still Exhibit Stress While ATM and ATR are Inhibited

DDR-induced cell apoptosis in response to Pt-DNA adducts is thought to act primarily downstream of ATM and ATR. Based on our H2AX and ATM assays, nucleolar stress inducing Pt(II) compounds did not appear to be significantly activating key initiation proteins in the DDR pathways associated with Pt-DNA repair. To further confirm these results, as well as investigate the role of ATR in Pt(II)-induced nucleolar stress, we next wanted to determine if nucleolar stress would still occur while inhibiting the primary DDR initiation proteins ATM and ATR.

Nucleolar stress can be measured by redistribution of the nucleolar protein NPM1 and quantified by coefficient of variation (CV) in cell samples (5, 37). NPM1 redistribution was measured after treatment with Pt(II) compounds in the presence of either ATM inhibitor (ATMi) (KU-55933) or ATR inhibitor (ATRi) (AZD6738), and compared to the NPM1 redistribution caused by the compounds without inhibitor. These studies were done at 3, 5, and 24-hour drug treatment times in A549, U-2 OS and HCT116 cell lines. For all three cell lines, nucleolar stress still occurred after treatment with ActD, OxaliPt, DACH-Pt and Penta-Pt in the presence of both ATMi and ATRi. Likewise, CisPt did not exhibit nucleolar stress in any of the cell lines in the presence or absence of ATMi and ATRi (Figure 5 **and S3-5**).

**Figure 5:**
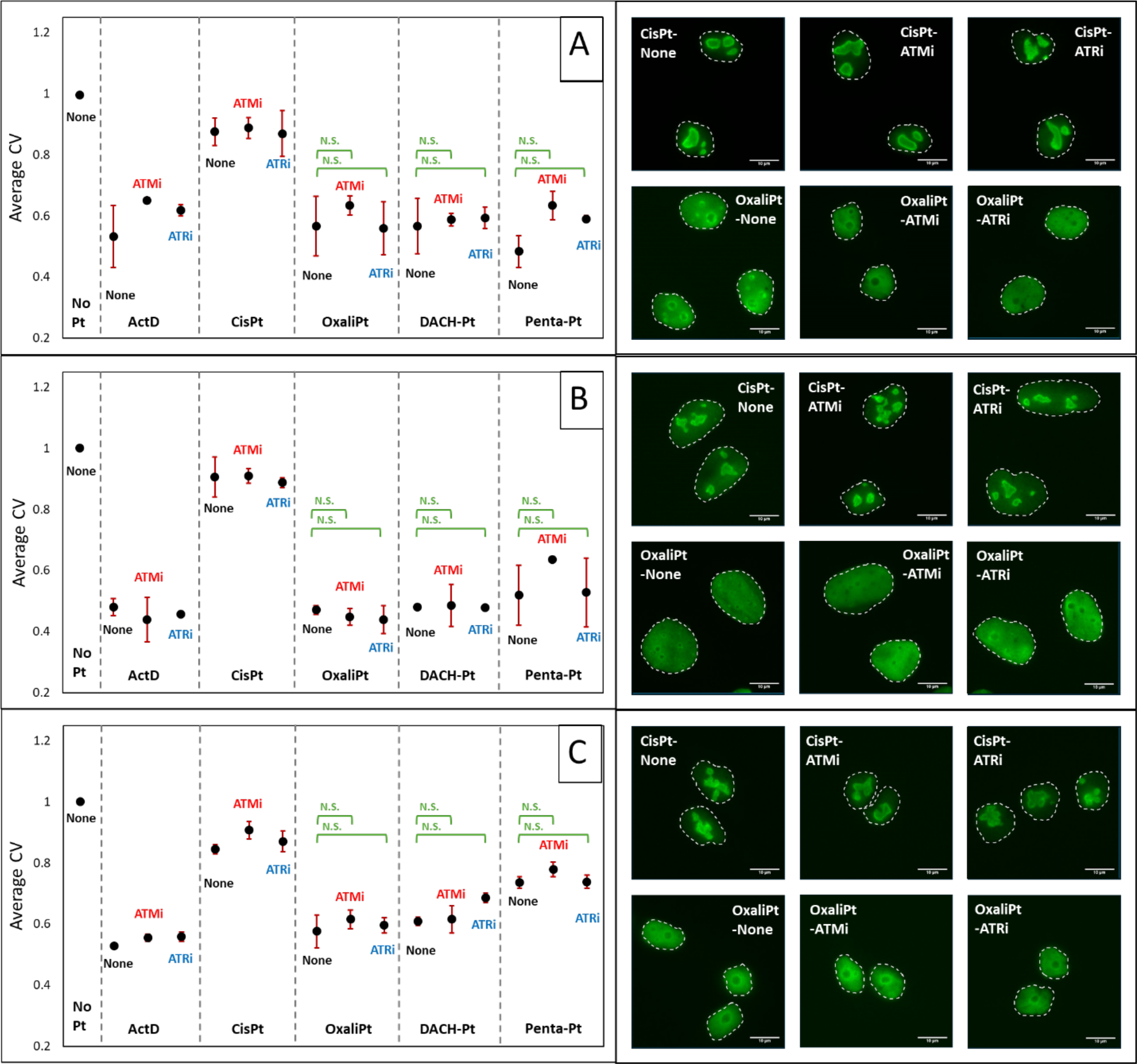
N**P**M1 **relocalization studies with ATM/ATR inhibitors at 24 hr. drug treatment.** Cells were treated with 10 µM platinum compounds or 5 nM of ActD for 24 hours in the presence or absence ATMi (10 µM) and ATRi (2.5 µM) in A549 (**A**), U-2 OS (**B**), and HCT116 (**C**) cell lines. NPM1 immunofluorescence distribution was then quantified (Methods). Each point is the average CV value and standard deviation for 3 biological replicates. Representative cell images of cells treated with **1** and **2** in the presence of ATMi, ATRi, or no inhibitor for each cell line are shown with NPM1 (green).

In A549 cells (Figure 5A), a slight decrease in the degree of NPM1 relocalization, indicated by a higher CV value, was observed for cells treated with ATM inhibitor at all three time points (results for the 3 and 5 hr treatment times can be found in **Figure S3)**. However, at each time point, this observed change was minimal, and all of the stress inducing compounds still exhibiting robust levels of NPM1 relocalization with ATMi, based on the cell images and the CV values ≈ 0.6 (Figure 5A **and S3**). Inhibition of ATR did not result in any significant changes to the degree of NPM1 relocalization at any of the 3 times points. In U-2 OS cells, we observe little to no influence on the degree of NPM1 redistribution when inhibiting ATM or ATR at any of the given time points (Figure 5B and **S4)**. Interestingly, we found that Penta-Pt takes longer to induce nucleolar stress in U-2 OS cells, regardless of treatment with ATM or ATR inhibitors, in comparison with A549 and HCT116 cells. This result mimics observations in the H2AX studies where U-2 OS cells showed a slower onset of activation, and supports the claim that Pt(II) compounds may initiate a slower response in U-2 OS cells possibly linked to the slower doubling time for this cell line. HCT116 cells (Figure 5C **and S5**) showed some variation between CV values with the addition of the inhibitors, but all of the nucleolar stress inducing Pt(II) compounds elicited pronounced NPM1 relocalization in the presence and absence of ATM and ATR inhibitors. Together these results suggest that neither ATR or ATM activation is essential for Pt(II)-induced NPM1 relocalization across various cell lines.

#### rRNA transcription is further decreased by treatment with OxaliPt when ATM and ATR are inhibited

Nucleolar stress caused by Pt(II) compounds is also characterized by loss of rRNA transcription (4, 5, 38, 70). We wanted to further investigate the roles of ATM and ATR in Pt(II)-induced nucleolar stress by measuring rRNA transcription levels after treatment with CisPt, OxaliPt, and ActD, in the presence and absence of ATM and ATR inhibitors. To perform these studies, quantitative PCR (qPCR) was used to determine relative rRNA transcription levels, based on the gene expression of the 5’ external transcribed spacer (ETS) region of pre-rRNA. The 5’ ETS region is commonly used to quantify relative rRNA transcription levels (43, 71, 72). Drug treatments were done at 24 hours in the presence or absence of inhibitor in A549 cells (Figure 6).

**Figure 6:**
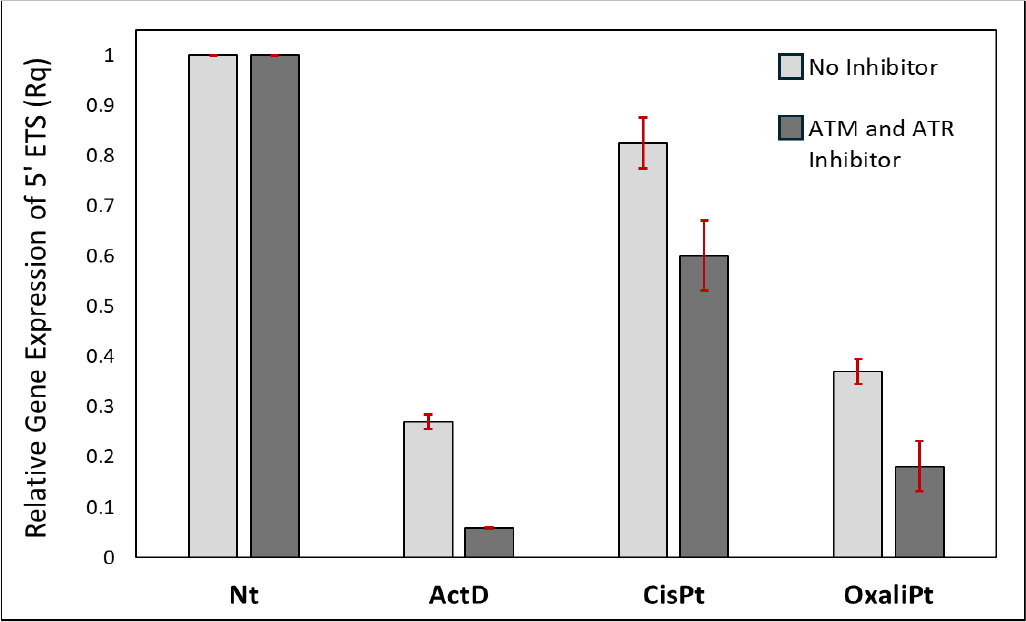
rRNA transcription qPCR studies with ATM and ATR inhibitor. A549 cells with inhibitors were treated with 10 µM ATMi and 2.5 µM ATRi for 12 hrs. prior to drug treatment. All cells were treated with 10 µM platinum compound or 5 nM ActD for 24 hrs. The relative expression of 5’ ETS was then determined (Methods) and plotted as an average with standard deviations from 3 biological replicates for cells with and without ATMi and ATRi.

Both OxaliPt and ActD treatments caused significantly decreased relative levels of rRNA transcription at 24 hours while CisPt only showed a minimal decrease in relative transcription levels (Figure 6) (5, 43). Interestingly, for all three compounds, the addition of ATM and ATR inhibitors resulted in a decrease in relative rRNA transcription levels.

These results indicate that ATM and ATR are not required for ribosomal biogenesis stress induced by Pt(II) compounds, and that inhibiting these DDR proteins actually makes the Pt(II) compounds more robust rRNA synthesis inhibitors. The decrease in relative transcription caused by CisPt in the presence of inhibitor may be attributed to DNA damage repair pathways being unable function properly in the absence of ATM/ATR, rendering higher levels of CisPt-induced DNA damage. This unrepaired DNA damage could possibly interfere with the transcription of rRNA from rDNA, or with other nucleolar processes (73, 74). Importantly, it should be noted that the decrease in relative transcription levels between treatments without inhibitor and those with inhibitor, is relatively similar for both CisPt and OxaliPt, indicating that there may still be some influence of DNA damage caused by OxaliPt on rRNA synthesis. OxaliPt treatment shows significantly lower rRNA transcription levels compared to CisPt under both conditions of no inhibitor and ATMi/ATRi treatments, however. These results, along with those from the ATMi/ATRi NPM1 relocalization studies, strongly suggest that Pt(II)-induced nucleolar stress does not require the activation of ATM or ATR.

### Pt(II)-induced nucleolar stress may involve G1 cell cycle arrest

To further investigate other potential mechanisms involved in Pt(II)-induced stress, we next explored potential correlations with cell cycle regulation and cell cycle arrest. In addition to being the location of ribosome biogenesis, another important role of the nucleolus is its impact on regulating cellular homeostasis. The nucleolus contains over 4,000 unique proteins, many of which are involved in sensing and initiating responses to various stimuli that impair cellular homeostasis (74). If cells are not able to recover from such stimuli, they will undergo cell cycle arrest or apoptosis, both of which outcomes can be regulated by nucleolar proteins(74). This regulation can be initiated through mechanisms by which displaced nucleolar proteins directly bind to other proteins that activate cell cycle arrest or apoptosis, or else sequester negative cell cycle regulators, leading to arrest or apoptosis. Generally, both mechanisms converge on the activation of tumor suppressor protein p53, which has various roles in cell cycle arrest, senescence, and apoptosis (75, 76). Given the important role the nucleolus plays in cell cycle regulation, we wanted to further investigate how cell cycle arrest at different phases may correlate with Pt(II)-induced nucleolar stress.

CisPt and OxaliPt induce different cell cycle arrest profiles across various cancer cell lines (29, 77, 78). Both compounds cause a relative increase in the number of cells in the G2 phase, however this increase is larger after CisPt treatment compared to OxaliPt (77). Key differences between the two compounds appear in the G1 and S phases, where CisPt treatment leads to a large decrease in the number of cells in the G1 phase and an increase of cells in S phase, whereas OxaliPt causes minimal changes to the number of cells in the G1 phase, while decreasing the number of cells in S phase (77, 79). Based on these findings it has been concluded that OxaliPt causes G1 cell cycle arrest while cisplatin causes S and G2 cell cycle arrest (77, 80). In addition to cell cycle analysis studies, gene expression profiles with OxaliPt treatment show key differences in the expression of cell cycle and proliferation proteins when compared to CisPt. Most notably, unlike CisPt, OxaliPt treatment causes downregulation of E2F, a transcription factor protein that is essential for the progression of cells into the S phase. This may be one of the key components involved in OxaliPt- induced G1 cell cycle arrest (29, 77). Based on these findings from previous studies, we wondered if the G1 cell cycle arrest induced by OxaliPt is an important factor in Pt(II)-induced nucleolar stress. To further investigate this, we measured key indicators of nucleolar stress caused by platinum compounds while inhibiting cell cycle at the G1-S and G2-M checkpoints.

### CisPt causes nucleolar stress with G1 cell cycle arrest

To investigate the correlation between Pt(II)-induced nucleolar stress and cell cycle arrest, we first aimed to observe NPM1 relocalization caused by Pt(II) compounds when cells were arrested in the G1 or G2 cell cycle phase. For these studies, A549 and U-2 OS cells were first treated with either Chk1 inhibitor (Caffeine, (81)) or Chk2 inhibitor (BML-277, (82)) for 12 hours prior to drug treatment, and then OxaliPt, CisPt or ActD were added in addition to the inhibitors for 3, 5 or 24 hours. Following drug treatment, the relocation of NPM1 was monitored (Figure 7). Interestingly, it was found that in the presence of Chk1 inhibitor, but not Chk2 inhibitor, CisPt caused robust relocalization of NPM1 in both cell lines by 24 hours of drug treatment. In A549 cells, with Chk1 inhibition, cisplatin-treated cells show NPM1 relocalization as early as 5 hours after drug treatment and robust NPM1 relocalization by 24 hours, indicated by the CV value of ≈0.6 and the corresponding cell images (Figure 7A **and S6**). This relocalization was not observed for cells treated with Chk2 inhibitor or cells treated with CisPt without the addition of inhibitor. OxaliPt and ActD caused robust NPM1 relocalization in the presence or absence of Chk1 and Chk2 inhibitors at all time points. Importantly, there was little to no NPM1 relocalization observed in cells that had been treated with Chk1 inhibitor, but no Pt(II) drug, indicating that it was not simply the inhibition of Chk1 that was causing the observed NPM1 relocalization.

**Figure 7:**
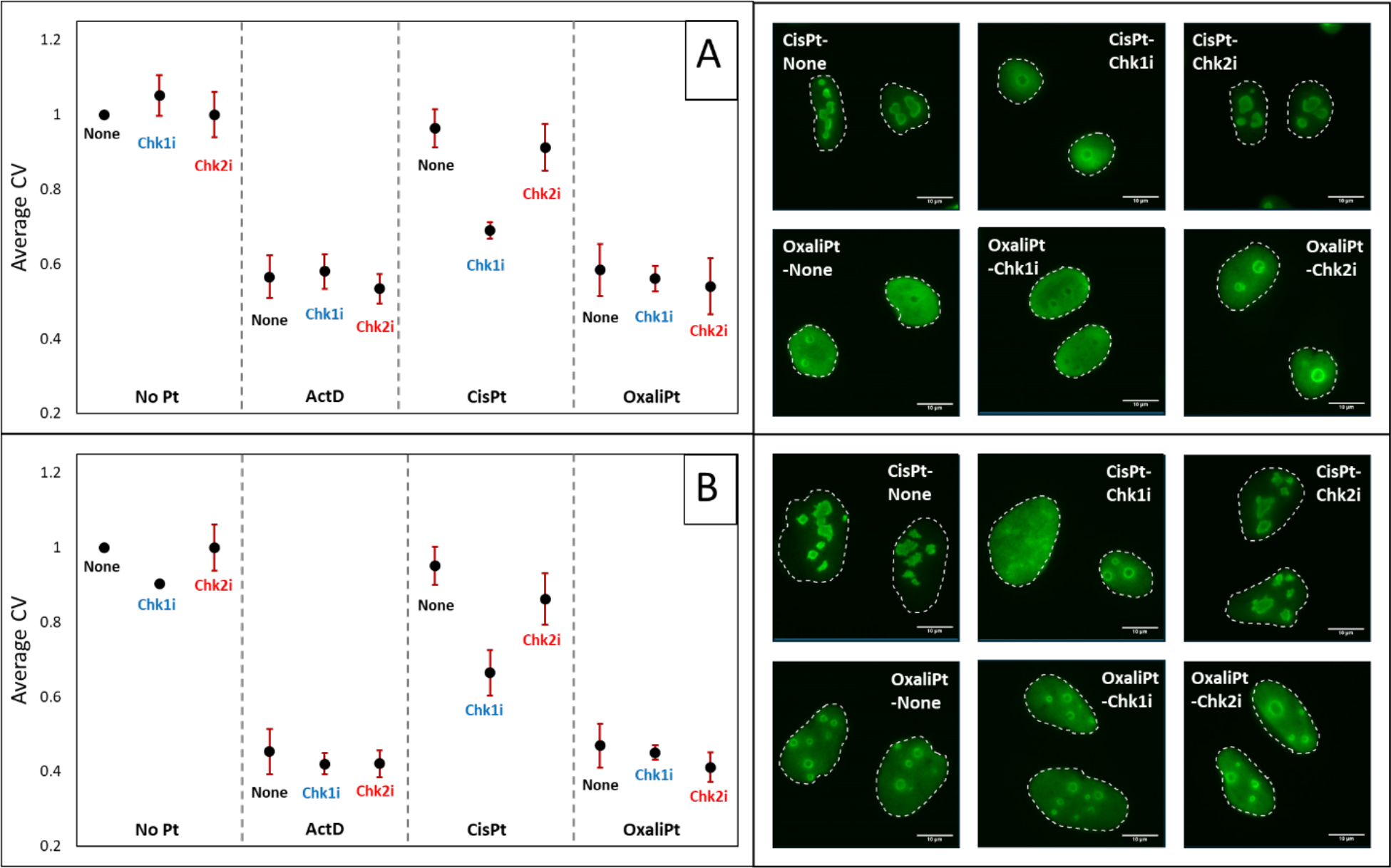
NPM1 redistribution with Chk1 and Chk2 inhibition at 24 hr. drug treatment. Cells were treated with 10 µM platinum compounds or 5 nM of ActD for 24 hours in the presence or absence of Chk1i (1.0 mM) or Chk2i (0.4 µM) in A549 (**A**) or U-2 OS (B) cell lines. NPM1 immunofluorescence distribution was then quantified (Methods). Each point is the average CV value and standard deviation for 3 biological replicates. Representative cell images of cells treated with 1 and 2 in the presence of Chk1i, Chk2i, or no inhibitor for each cell line are shown with NPM1 (green).

U-2 OS cells (Figure 7B) show similar results to those of A549 cells, where by 24 hours CisPt as well as OxaliPt and ActD induce robust NPM1 relocalization in the presence of Chk1 inhibitor, while little change is observed with Chk2 inhibitor compared to the drug treatments in the absence of inhibitor. Compared to A549 cells, CisPt induced relocalization with Chk1 inhibited occurred slower in U-2 OS cells, with only slight relocalization observed at the 5-hour drug treatment time (**Figure S7**). This may be a further indication of the slower onset of activation of platinum compounds described above for U-2 OS cells.

### CisPt causes fibrillarin cap formation when cells are arrest in the G1 phase

To further confirm the results of the NPM1 relocation studies with Chk1 inhibition, we monitored another key indicator of nucleolar stress, fibrillarin cap formation. Upon nucleolar stress induction, key proteins in the nucleolus reorganize and form distinct morphologies when compared to cells not undergoing stress.

Of these proteins, fibrillarin has commonly been utilized to monitor nucleolar stress induction, as it forms nucleolar caps along the periphery of nucleoli upon stress-induction (83, 84). Previous studies from our lab have shown that at 24 hour drug treatment times, OxaliPt and derivatives cause the formation of fibrillarin caps, while CisPt treatment shows little cap formation (38, 40, 41). Given the robust NPM1 relocalization observed for CisPt with Chk1 inhibition, we were interested in determining if fibrillarin cap formation was also occurring under these conditions.

In these studies, fibrillarin immunofluorescence in the presence and absence of Chk1 inhibition was monitored in A549 (Figure 8A) and U-2 OS (Figure 8B) cells. Cells were treated with Chk1 inhibitor for 12 hours prior to drug treatment, and then OxaliPt, CisPt or ActD were added for an additional 5 and 24 hour treatment periods (Figure 8 **and S8).** In the absence of platinum compounds fibrillarin distributes evenly in the nucleolus, with and without Chk1 inhibitors. OxaliPt and ActD caused clear cap formation in the presence and absence of Chk1 inhibitor at 5 and 24 hours drug treatments. In the absence of Chk1 inhibitor, cisplatin treatment does not cause fibrillarin cap formation. With addition of Chk1 inhibitor, however, cisplatin-treated A549 cells show fibrillarin cap formation as early as 5 hours after drug treatment (**Figure S8**). By 24 hours of drug treatment, nearly all cisplatin-treated cells that were observed showed the formation of pronounced fibrillarin caps at the periphery of the nucleoli (Figure 8A).

**Figure 8:**
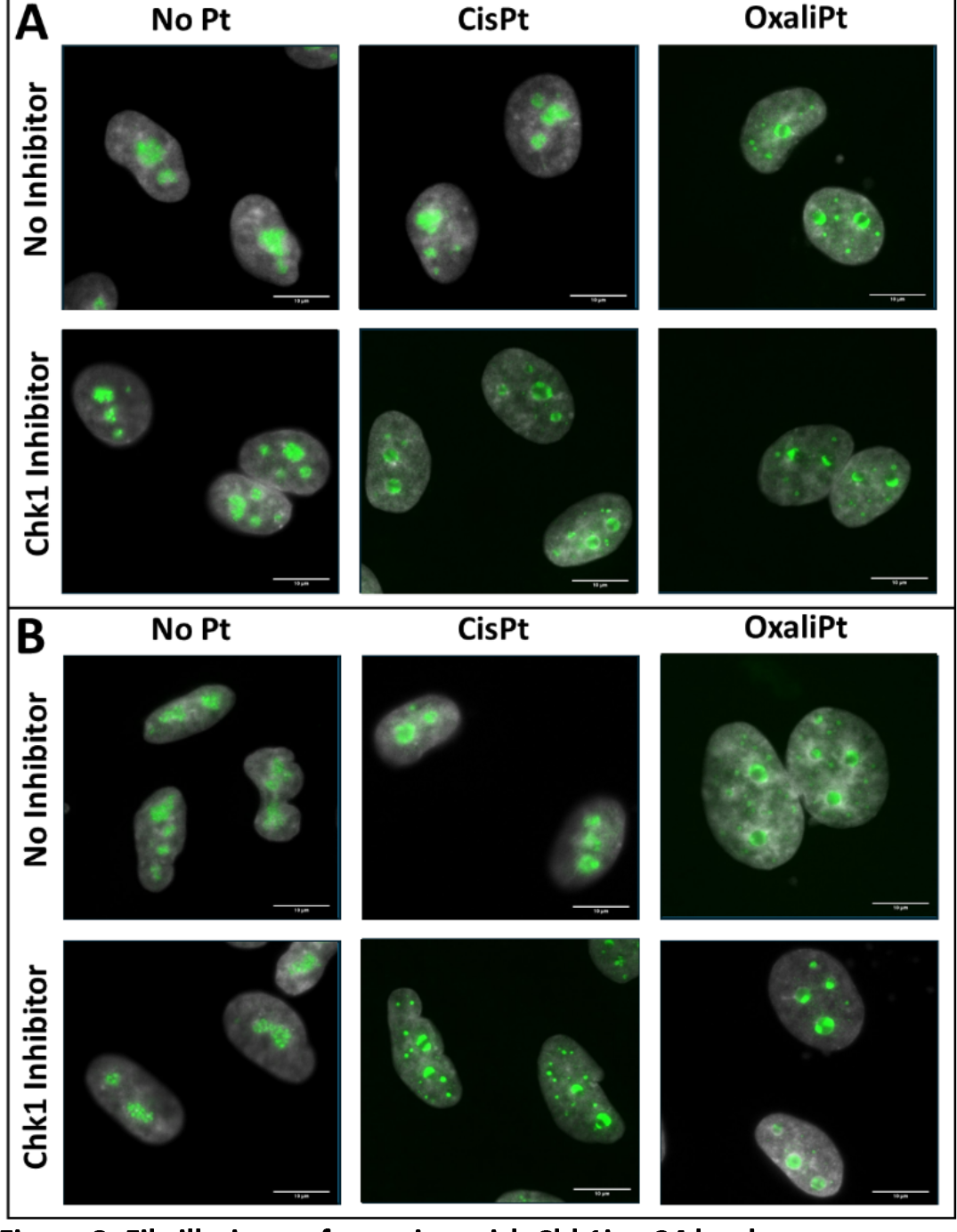
F**i**brillarin **cap formation with Chk1i at 24 hr. drug treatment**. Cells were treated with 10 µM platinum compounds or 5 nM of ActD for 24 hours in the presence or absence of Chk1i (1.0 mM) in A549 (**A**) or U-2 OS (**B**) cell lines. Fibrillarin cap formation was then determine based on immunofluorescent images. Representative images for the no drug control and cells treated with **1** and **2** in the presence and absence of Chk1i are shown with fibrillarin (green) overlayed with DAPI (grey).

U-2 OS cells (Figure 8B **and S8**) showed results similar to A549 cells. While fibrillarin cap formation upon CisPt treatment in the presence of Chk1i was less pronounced at the shorter treatment times (**Figure S8**), matching the timeframe of NPM1 relocalization in these cells (**Figure S7**), by 24 hr drug treatment, pronounced fibrillarin cap formation was observed (Figure 8B).

Together these results further confirm observations from NPM1 relocalization studies that CisPt appears to be inducing robust nucleolar stress when cell cycle arrest is halted at the G1-S checkpoint in A549 and U-2 OS cells at 24 hour drug treatment time. As with NPM1 redistribution, fibrillarin cap formation has a slower onset for CisPt treatment compared to OxaliPt, further indicating that OxaliPt may be eliciting its nucleolar stress effects through other interactions in addition to G1 cell cycle arrest.

### G1 cell cycle arrest increases Pt(II)-induced RNA transcription inhibition

In a final experiment aimed at observing nucleolar stress induction following G1 cell cycle arrest, RNA transcription levels were measured following Chk1 inhibition and drug treatment. In these studies, A549 cells were treated with Chk1 inhibitor for 12 hours prior to drug treatment, and then OxaliPt, CisPt or ActD were added for an additional 24-hour treatment period. To measure newly synthesized RNA, 5-ethynl uridine (5-EU) was incorporated into cells during the final 4 hours of drug treatment. Cells were fixed and then labeled with the alkyne containing Oregon green 488 dye using Cu(I)-catalyzed azide–alkyne 1,3- dipolar cycloaddition (CuAAC) (85). The fluorescent intensity of 5-EU within the cellular nucleus, segmented by DAPI stain, was then measured, and used to quantify RNA transcription levels (Figure 9).

**Figure 9:**
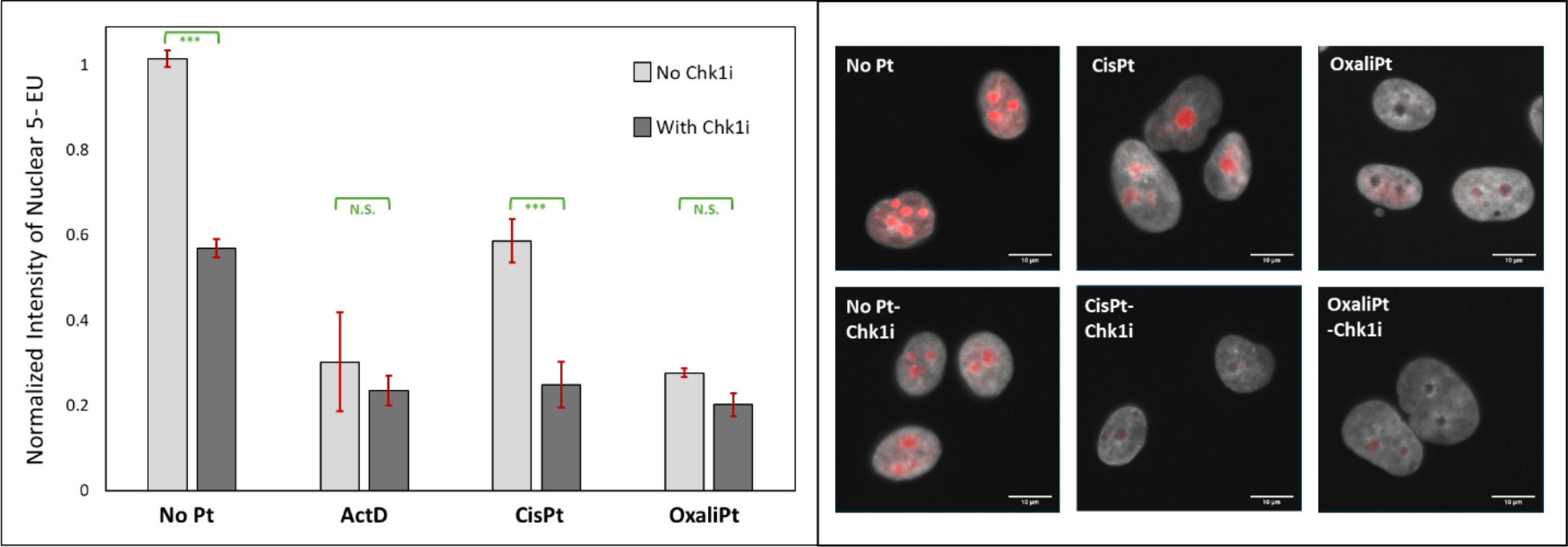
R**N**A **transcription inhibition studies with Chk1i at 24 hour drug treatments with 5-EU quantification**: “Inhibited” cells were treated with Chk1i (1mM) for 12 hours prior to drug treatment. All cells were treated with 10 µM platinum compounds or 5 nM of ActD for 24 hours. During the final 4 hours of drug treatment, 5-EU (3mM) was incorporated into newly synthesized RNA. Following treatment, 5-EU was labeled with alkyne-containing Oregon green 488 dye, using CuAAC. The intensity of 5-EU was then measured within the nuclei of cells, segmented by DAPI, and normalized to that of the no drug treatment control. The above graph shows the average and standard deviations from 3 biological replicates. Representative cell images for the no treatment control, **1** and **2** in the presence and absence of Chk1i, are shown with 5-EU (red) and DAPI (grey).

In the absence of Chk1i, OxaliPt and ActD treatments both cause significant inhibition of RNA transcription, showing ∼80% decrease in transcription compared to that of the no drug control, while CisPt treatment only produces around a ∼40% decrease (Figure 9) (5, 38). Treatment with Chk1i causes a decrease in transcription for all cells. This is expected, as DNA replication takes place in S phase and cells halted in the G1 cell cycle phase have overall lower levels of DNA available for RNA transcription (42). Chk1i treatment has little effect on already-reduced transcription levels for OxaliPt and ActD samples. Pre-treatment of cisplatin-treated cells with Chk1i, however, causes a further significant decrease in transcription levels, with an additional ∼40% decrease in transcription compared to the no Pt(II) control.

Together these results, combined with those observed for NPM1 relocalization and fibrillarin capping with Chk1 inhibition, show that cisplatin treatment can induce robust nucleolar stress when cells are blocked from entering G1, whereas nucleolar stress induced by OxaliPt is unaffected by Chk inhibitors. The results from these studies support two important conclusions related to Pt(II)-induced nucleolar stress. First, cells can undergo nucleolar stress in the absence of entering the S and G2 phase, which further confirms that DDR, which occurs primarily in the S phase (86), is not necessary for nucleolar stress to occur. Second, when cells are arrested in the G1 phase, CisPt treatment can cause nucleolar stress, indicating that G1 cell cycle arrest may be a pivotal factor in Pt(II)-induced stress, and may be closely related to the ability of OxaliPt, but not CisPt, to cause stress in cells without inhibitor. However, based on the observation that CisPt still takes longer to induce nucleolar stress in the presence of Chk1i compared to OxaliPt, it can be speculated that cell cycle arrest at the G1 checkpoint is not the sole contributor to the ability of Pt(II) compounds to induce stress. There are likely other important interactions occurring with OxaliPt, but not CisPt, that elicit its more profound nucleolar stress-inducing capabilities.

### Pt(II) induced nucleolar stress is non-reversible and unique from other small molecule nucleolar stress inducing compounds

#### Nucleolar stress inducing compounds Act D, BMH-21 and CX-5461 cause reversible nucleolar stress

To further understand mechanisms by which some platinum compounds cause nucleolar stress, we compared properties with other molecules known to inhibit rRNA transcription and induce nucleolar stress. ActD, BMH-21 and CX-5461 are all small organic molecules that have been shown to induce nucleolar stress in various cell lines (87–89). The mechanisms of actions for the three compounds, although extensively studied, have still not been fully elucidated and may be cell line dependent (90–92). ActD is thought to act as a DNA intercalator, interacting with GC-rich regions of DNA and preventing transcription of RNA, DNA, and protein synthesis (33, 93); at low concentrations, ActD is selective for rRNA synthesis inhibition (94). The primary mechanism for BMH-21 is to also act as a DNA intercalator, interacting at GC-rich regions of DNA and preventing transcription. Interestingly, BMH-21 has been shown to have a specific influence on RNA polymerase I (Pol I) and in addition to inhibiting RNA Pol I from binding to rDNA to activate rRNA transcription, it has also been shown to cause proteasome-dependent destruction of RPA194, the large catalytic subunit of RNA Pol I (95–97). The mechanism for CX-5461 is highly debated in the field and has been attributed to being a topoisomerase II poison, a p53-mediated DDR activator, and has also been shown to inhibit rRNA transcription by arresting the transcription initiation complex for RNA Pol I (98–102). Oxaliplatin and DACH-Pt are chemically distinct from these organic molecules but are also expected to form stable complexes with cellular targets in order to elicit their effects. We wondered if oxaliplatin and other stress-inducing Pt(II) compounds were working through mechanisms similar to that of the ActD, BMH-21 or CX-5461. In order to test this, we measured the reversibility of NPM1 relocalization following treatment with ActD, BMH-21, CX-5461, and nucleolar stress-inducing Pt(II) compounds. ActD and BMH-21 have both been shown to cause reversible rRNA transcription inhibition, in which cells can recover after drug removal and can restore rRNA transcription (89, 103). CX-5461 however, has previously been shown to cause irreversible rRNA transcription inhibition and nucleolar stress (102, 104).

We initially wanted to confirm the reversibility of nucleolar stress induced by ActD, BMH-21, and CX-5461 in A549 cells. For these studies, we first needed to determine what drug treatment concentration to use for each compound. We commonly use ActD as a positive control for nucleolar stress at the clinically relevant concentration of 5 nM. For BMH-21, previous work focused on rRNA transcription inhibition with this compound and utilized concentrations between 1-5 µM, thus a 3 µM concentration was chosen for our reversibility studies. For CX-5461, a larger variation of concentrations has been used in previous studies, ranging from 0.05-5.0 µM. We sought to determine clinically relevant concentrations that were able to induce nucleolar stress for this compound in A549 cells, prior to running reversibility studies. We tested a range of concentrations for CX-5461 that were slightly above and below the previous report IC50 value in A549 cells of 0.169 (-/+3.74) µM (**Figure S9**) (88). Cells were treated for 3 hours with CX-5461 and nucleolar stress was measured based on NPM1 relocalization. From these results, we chose the treatment concentrations of 0.2 µM to be utilized in the reversibility studies. We next looked at the reversibility of NPM1 relocalization after 3 hour drug treatment times followed by 24 hour drug free media recovery periods for ActD, BMH-21, and CX-5461 (**Figure 10A**). As expected, we found the NPM1 relocalization for ActD and BMH-21 was reversible after the 24-hour recovery step. Interestingly, we found that CX-5461 at 0.2 µM treatment concentration is also reversible after the recovery step (**Figure 10B**). Of note, CX-5461 concentrations used in prior experiments showing irreversible behavior are significantly higher (1 µM-10 µM) than the 0.2 µM concentration utilized in these studies and may be responsible for the observed differences in reversibility (101, 102). To test this, NPM1 assays were utilized for cells treated with CX-5461 at 0.05- 2.0 µM concentrations for 3 hours, followed by a 24 hour drug-free recovery period, and indeed demonstrated that drug treatment at concentrations of 0.5 µM or higher, render nucleolar stress by CX- 5461 irreversible, while treatments at 0.2 µM or lower induce reversible NPM1 relocalization (**Figure 10B**).

**Figure 10:**
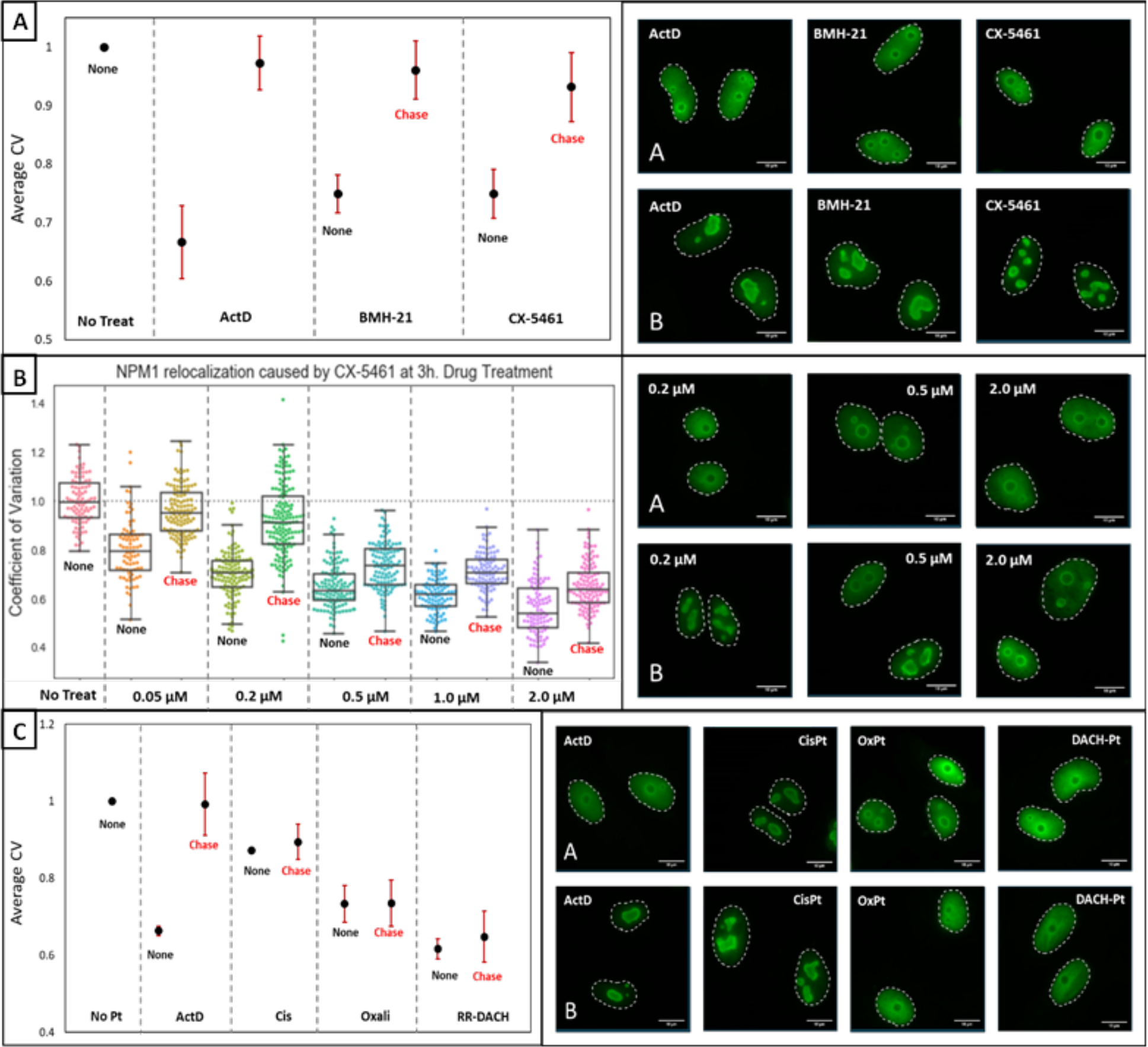
N**P**M1 **relocalization with small molecule nucleolar stress inducing compounds.** A549 cells were treated with ActD (5 nM), CX-5461 (0.2 µM), and BMH-21 (3.0 µM) (**A**), various concentrations of CX-5461 (**B**), or with ActD (5 nM) and 10 µM platinum compounds (**C**) for 3 hours. The drug was removed, and drug free media was replaced for 24 hr. NPM1 immunofluorescence distribution was then quantified (Methods). For (**A** and **C**) each point is the average CV value and standard deviation for 3 biological replicates. Representative cell images of NPM1 (green) are shown after 3 hr. drug treatment alone (**A**), and 3 hr. drug treatment followed by 24 hr. drug free chase (**B**).

We also tested the reversibility of NPM1 relocalization caused by Pt(II) compounds. Unlike the other small molecule inhibitors, all of the nucleolar stress inducing Pt(II) compounds showed irreversible NPM1 relocalization following 3-hour drug treatment and 24-hour drug-free recovery (**Figure 10C)**. Similarly, rRNA transcription as measured by qPCR also shows irreversible inhibition by oxaliplatin treatment, whereas inhibition by ActD is reversed (**Figure S10**). The results from these studies may indicate that Pt(II) compounds are inducing nucleolar stress through a unique mechanism, unlike that of ActD, CX-5461, or BMH-21. All of the small organic molecule inhibitors have been show to at least in part act as intercalators, while Pt(II) compounds cause primarily bi-functional metal-biomolecule binding (105). These results suggest that if the Pt(II) compounds are causing nucleolar stress by binding to nucleic acids in RNA, DNA, or proteins, that binding and its effects may be irreversible.

## Conclusions

The overarching goal of this work was to better understand the unique nucleolar stress pathway induced by some Pt(II) compounds, and how it relates to the canonical DNA Damage Response. Since ribosome biogenesis occurs in the nucleolus of cells, it has been speculated that oxaliplatin may elicit its effects through the nucleolar-DNA damage response (n-DDR) pathway (42). n-DDR primarily occurs in cells that have accumulated DNA damage on rDNA, which is housed primarily in the nucleolus. It is thought that upon nucleolar DNA damage, ATM or ATR will recognize the damage and initiate a protein signaling cascade, leading to the regulation of cell cycle progression, DNA replication and RNA transcription. Specifically, upon recognition of rDNA DSBs by ATM, rRNA transcription via Pol I is inhibited and damaged rDNA migrates into nucleolar caps where it can then be repaired by various DNA repair proteins that are segregated to the nucleolar caps upon ATM activation. One such ATM-activated nucleolar protein, Treacle (TCOF1), can recruit DNA topoisomerase II binding protein 1 (TOPBP1) to activate ATR down stream of ATM activation, leading to the complete shutdown of rRNA transcription (73, 74, 106, 107). Since Pt(II) compounds are known to cause DNA DSBs during DNA repair and replication, we considered if n-DDR was possibly involved in rRNA transcription inhibition elicited by oxaliplatin and derivatives.

In agreement with recent findings from Kleiner and colleagues, results from this manuscript strongly suggest the n-DDR is not likely the main driver of ribosome biogenesis inhibition by oxaliplatin (42) and moreover that n-DDR may even hinder the compound from inducing its full transcriptional inhibitory effects. The finding that oxaliplatin and derivatives cause low activation of key DDR proteins compared to cisplatin, specifically ATM activation (Figure 4), suggests little involvement of ATM-activated n-DDR in oxaliplatin’s mechanism. One may question if the lower degree of ATM activation is simply a product of the lower DNA adduct formation by oxaliplatin compared to cisplatin. However, the small amount of ATM activation induced by oxaliplatin treatment can be reversed by 24 hr incubation in Pt-free media, (**Figure S2**), whereas nucleolar stress is still occurring after this recovery period (Figure 10**, S10**). By contrast, cisplatin-treated cells still show predominant activation of ATM and H2AX after drug-free recovery, while still not inducing rRNA transcription inhibition. Other ATM and ATR inhibition studies also support the lack of involvement of n-DDR in oxaliplatin’s cell death mechanism. Specifically, inhibiting ATM or ATR has little to no effect on the ability of oxaliplatin, and other stress-inducing compounds, to cause NPM1 relocalization (Figure 5). Even more significantly, we see that inhibiting ATM and ATR not only fails to relieve inhibition of rRNA transcription by oxaliplatin, but that transcription levels decrease even further with the addition of ATM/ATR inhibitor compared to oxaliplatin treatment alone (Figure 6). This strongly suggests that RNA transcription inhibition in response to oxaliplatin treatment is not mediated by ATM, and moreover that any ATM-mediated n-DDR occurring in response to oxaliplatin treatment may be impeding compound’s ability to induce ribosome biogenesis stress. This observation may be relevant to studies that have shown that various cancer lines become more sensitive to oxaliplatin treatment when ATM is inhibited, and may also be related to why oxaliplatin is most effective in treating colorectal cancers, which are often ATM deficient (46, 108, 109).

Since the main regulators of DDR do not appear to play a significant role in oxaliplatin-induced nucleolar stress or cell death, we next investigated other potential cellular response pathways that may be involved in oxaliplatin’s mechanism. Given the significant role that the nucleolus plays in regulating cell cycle progression and cell division, we wanted to determine potential links between cell cycle arrest and Pt(II)- induced nucleolar stress. Interestingly, we found that there may be a significant link between G1 cell cycle arrest and platinum-induced nucleolar stress. Treatment with a Chk1 inhibitor, inhibiting the G1-S transition, does not influence the ability of oxaliplatin and related derivatives to cause nucleolar stress. However, surprisingly, we observe that CisPt causes substantial nucleolar stress upon G1 cell cycle inhibition (Figure 7**-9**). This observation may indicate that there is some molecular interaction or adduct formation occurring with oxaliplatin in the G1 phase that allows it to elicit its primary nucleolar effects, and that cisplatin may also be capable of undergoing this molecular interaction, but with influences on the nucleolus only when cells are unable to proceed through the cell cycle. This may be because DDR primarily occurs in the S-Phase and under normal cell division, may prevent this interaction from occurring for cisplatin, but not oxaliplatin, through use of DDR repair pathways. However, compared to our rRNA transcription studies with ATM/ATR inhibition (Figure 7), there is a significantly larger decrease in rRNA transcription from cisplatin with Chk1 inhibition, indicating that cisplatin induced inhibition may not be ATM/ATR dependent, further supporting the claim that Pt(II)-induced stress is not working through n-DDR. Future studies are needed to further explore potential molecular interactions involved in this cell cycle dependent stress response, as there are multiple cellular pathways involved with G1 cell cycle arrest and cell cycle regulation.

To further investigate potential mechanisms for nucleolar stress induction by oxaliplatin and derivatives, we performed comparison studies with other small-molecule rRNA transcription inhibitors, ActD, BMH- 21, and CX-5461. While 3-hour treatment with each of these different organic small molecules causes robust nucleolar stress, the changes could be reversed on incubation in treatment-free media. By contrast, nucleolar stress induced by oxaliplatin and Pt(II) derivates was irreversible (Figure 10). An interesting case was rRNA transcription inhibitor CX-5461, which shows a gradient of reversibility across drug treatment concentrations. Specifically, at low treatment concentrations (0.05-0.3 µM) nucleolar stress induced by CX-5461 appears to be reversible, but at higher treatment concentrations (0.5-2.0 µM) the nucleolar stress is no longer reversible. This interesting finding suggests an initial and reversible binding event by CX-5461 that induces nucleolar stress morphologies, and an irreversible effect occurring at higher drug concentrations.

Taken together, these studies sharpen the contrast between DDR induction by cisplatin and nucleolar stress induction by oxaliplatin, and moreover strongly suggest that oxaliplatin does not act through a nucleolar-DDR mechanism. Many questions remain regarding how oxaliplatin is inducing its cellular effects. Specifically, one area of high importance is determining how oxaliplatin causes rRNA transcription inhibition. From these studies and others, it can be concluded that ATM-mediate n-DDR is not likely involved in transcription inhibition by oxaliplatin, allowing future studies to focus on other known pathways that cause rRNA transcription inhibition, and how they compare to platinum induced stress (110–113).

## Experimental Procedures

### Cell culture and treatment

A549 human lung carcinoma cells (#CCL-185, American Type Culture Collection), HCT116 human colorectal carcinoma cells (#CCL-247, American Type Culture Collection, and U-2 OS human osteosarcoma cells (#HTB-96, American Type Culture Collection) were cultured at 37°C, 5% CO2 in Dulbecco’s Modified Eagle Medium (DMEM) for A549 and HCT116 cells and Mccoy’s 5A Medium for U-2 OS cells that was supplemented with 10% Fetal Bovine Serum (FBS) and 1% antibiotic-antimycotic. Treatments were conducted on cells that had been grown for 11-30 passages to 70-80% confluency. Platinum compound treatments were conducted at 10 µM, Actinomycin D treatments were conducted at 5 nM, BMH-21 treatments were conducted at 3 µM and CX-5461 treatments were conducted at 0.2 µM unless otherwise noted. Actinomycin, BMH-21 and CX-5461 stocks were stored frozen in DMSO and thawed on day of use. Platinum compounds were made into 5mM stocks on the day of treatment from solids in water (oxaliplatin and cisplatin), or DMF (remaining platinum compounds). Stock solutions were diluted into media immediately prior to drug treatment. All inhibitor compounds were added to cells 12 hrs. prior to drug treatment and remained for the entirety of the treatment. ATM inhibitor (KU-55933) treatment was conducted at 10 µM, ATR inhibitor (AZD6738) treatment was conducted at 2.5 µM, Chk1 inhibitor (caffeine) treatment was conducted at 1 mM, and Chk2 inhibitor (BML-277) treatment was conducted at 0.4 µM. Ku-5593, AZD6738, and BML-277 stocks were stored frozen in DMSO and thawed on day of use. Caffeine stocks were made at 100 mM in water and heated to 95 C° to ensure solubility prior to use on day of treatment. For reversibility studies cells were washed 3X with PBS prior to the addition of free drug free medium.

### Immunofluorescence

Cells were grown on coverslips (Ted Pella product no. 260368, round glass coverslips, 10 mm diam. 0.16–0.19 mm thick) as described above. After treatment was complete, cells were washed with phosphate buffered saline (PBS) and fixed with 4 % paraformaldehyde (PFA) in PBS for 20 minutes at room temperature. PFA was removed using aspiration and cells were permeabilized with 0.5 % Triton-X in PBS for 20 minutes at room temperature. Two ten-minute blocking steps were then performed with 1 % bovine serum albumin (BSA) in PBST (PBS with 0.1 % Tween-20). Cells were incubated for one hour in primary antibody NPM1 (FC-61991, ThermoFisher, 1 : 500 dilution in PBST with 1 %BSA) , γH2AX (CR55T33, ThermoFisher, 2.5 μg in PBST with 1% BSA) , pATM (), or fibrillarin (ab4566 from Abcam, 1:500 dilution in PBST with 1% BSA), and then washed 3X with PBST and incubated for 1 hour in secondary antibody Goat Anti-Mouse IGG H&L Alexa Fluor® 488 (ab150113, Abcam, 1 : 1000 dilution in PBST with 1 % BSA) and washed again 3X in PBST prior to mounting the slides. Coverslips were then mounted on slides with ProLong™ Diamond Antifade Mountant with DAPI (ThermoFisher) according to manufacturer’s instructions.

### Image processing and quantification

Images were taken using a HC PL Fluotar 63x/1.3 oil objective mounted on a Leica DMi8 fluorescence microscope with Leica Application Suite X software. The quantification of NPM1 relocalization was performed in an automated fashion using a Python 3 script. Images were preprocessed in ImageJ,^36,37^ to convert the DAPI and NPM1 channels into separate 16-bit grayscale images. Between 50-250 cells were analyzed for each treatment group. Nuclei were segmented using the DAPI images using Li thresholding function in the Scikit-Image Python package.^38^ The coefficient of variation (CV) for individual nuclei, which is defined as the standard deviation in pixel intensity divided by the mean pixel intensity, was calculated from the NPM1 images using the SciPy Python package. All the data was normalized to the no-treatment in each experiment. NPM1 imaging results for each complex were observed in triplicate. Data are represented as boxplots generated using Seaborn within Python.

Quantification of γH2AX and pATM intensity and foci was performed with CellProfiler 4.2.1 software.^39^ In one analysis method, a “percent positive” value was calculated for each treatment condition relative to the untreated control. A threshold was determined for a positive γH2AX result based on the 90th percentile intensity value of the untreated control for each time point. Nuclei in the experimental samples with integrated intensity levels higher than this were counted as positive for γH2AX or pATM.

### 5-ethynyl uridine (5-EU) RNA labeling

Cells were grown on coverslips (Ted Pella product no. 260368, round glass coverslips, 10 mm diam. 0.16–0.19 mm thick) as described above. Cells treated with caffeine were incubated for twelve hours with caffeine before drug treatment. Cells were treated with the compound of interest with and without caffeine for a total of twenty-four hours. During the last four hours of treatment, RNA labeling was conducted. The cells were washed with PBS 3x at the twenty-hour mark, and media containing both the compound, with and without caffeine and 3 mM 5-EU were added to the cells and incubated for 4 h. Control treatments for no 5-EU had all components added excluding the 5-EU. Following treatment cells were washed with PBS 3X for 5 minutes each and fixed with 4% PFA in PBS for 20 minutes. Cells were then permeabilized with 0.5% Triton-X in PBS for twenty minutes. Block was performed with 5% BSA in PBST for one hour. During the last ten minutes of blocking, a click cocktail containing 10 mM sodium ascorbate, 5 μM Oregon Green™ 488 Azide (ThermoFisher, O10180), and premixed 2 mM CuSO4 and 4 mM THPTA (Lumiprobe, 760952-88-3) in PBS was made and then added to cells for one hour. Control treatments for no Cu had all components of the click cocktail added excluding the CuSO4. After the click reaction, cells were then washed with 0.5% Triton-X in PBS 1X for twenty minutes and in PBS 3X for five minutes each. Coverslips were mounted on slides with antifade fluorescence mounting media (Abcam) and left to cure overnight before imaging.

### qPCR RNA transcription quantification

Cells were grown in 100mm cell culture plates and treated as described above. Following treatment, total cellular RNA was extracted using Quick-RNA Purification Kit (Zymo, R1054). RNA was then treated with DNase I for 30 minutes according to manufacturer instructions (Thermo Fisher, EN0521) to remove any potential DNA contamination. Cellular RNA was then reverse transcribed to cDNA using Qscript cDNA Synthesis Kit (VWR, 101414-098). Two step qPCR was then ran using iQ SYBR Green supermix (BioRad, 1708880) and primers for the target gene, 5’ ETS (FP-CTTTCTCGCGCCTTCCCC; RP- GGGAGAAGACGAGAGACCAC) and the reference gene B2M (FP-AAGATGAGTATGCCTGCCGTGT; RP-TCTTCAAACCTCCATGATGCTGCT) at 300 nM primer and 75 ng of cDNA template. The following thermal cycling conditions were used: 1 cycle 95 °C (3 minutes), 40 cycles of 95 °C (10 seconds) and 60 °C (30 seconds). qPCR was run and results were analyzed using Quantum Studio 3 software. Relative gene expression (rq) was then calculated using the reference gene B2M and normalized to the no treatment controls. For each biological replicate, samples were run in triplicates and the average rq values for each sample were recorded with standard deviations < 0.2 between replicate samples. At least three biological replicates were run for each experiment across three different treatment days.

### Data Availability

Primary data are available upon request

## Supporting Information

This article contains Supporting Information.

## Acknowledgements

We would also like to recognize Danah Hijaz, Monique Demuth and Manasa Rajeev for their early contributions to the reversibility studies done in this manuscript.

## Funding and additional information

We gratefully acknowledge support from the NSF through CHE 2109255 (VJD) and DGE 2022168 (KRA).

