## Supplemental Information for "The Unique Pt(II)-Induced Nucleolar Stress Response and its Deviation from DNA Damage Response Pathways"

### Table of Contents

**Page Number**

Supplemental Figure 1 (**S1**) S2
**Quantification of H2AX activation after 48 hr. drug treatments**

Supplemental Figure 2 (**S2**) S2
**Quantification of ATM activation after 5 hr. drug treatment and a 24 hr drug free chase period**

Supplemental Figure 3 (**S3**) S3
**NPM1 relocalization studies with ATM/ATR inhibitors at 5 and 3 hr drug treatment in A549 cells**

Supplemental Figure 4 (**S4**) S4
**NPM1 relocalization studies with ATM/ATR inhibitors at 5 and 3 hr drug treatment in U-2 OS cells**

Supplemental Figure 5 (**S5**) S5
**NPM1 relocalization studies with ATM/ATR inhibitors at 5 and 3 hr drug treatment in HCT116 cells**

Supplemental Figure 6 (**S6**) S6
**NPM1 redistribution with Chk1 inhibition at 5 and 3 hr drug treatment in A549 cells**

Supplemental Figure 7 (**S7**) S7
**NPM1 redistribution with Chk1 inhibition at 5 and 3 hr drug treatment in U-2 OS cells**

Supplemental Figure 8 (**S8**) S8
**Fibrillarin cap formation with Chk1i at 5 hr drug treatment**

Supplemental Figure 9 (**S9**) S9
**NPM1 relocalization with small molecule nucleolar stress inducing compounds**

Supplemental Figure 10 (**S10**) S9
**rRNA transcription qPCR studies at 5hr. drug treatment and 24 hr. drug-free chase**

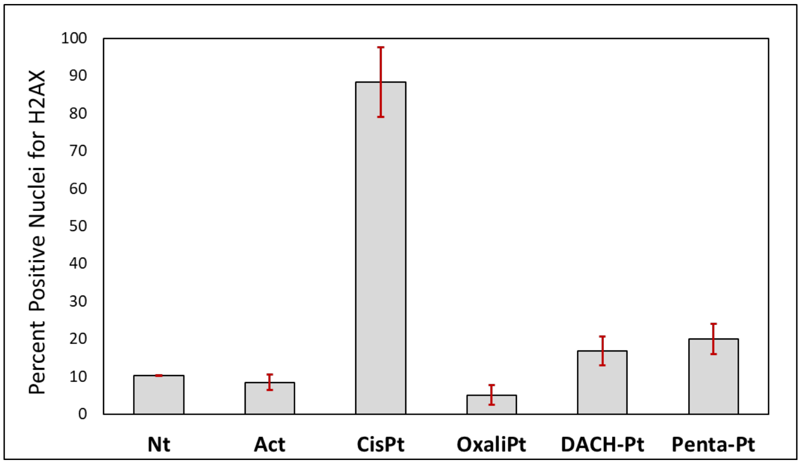


**Figure S1: Quantification of H2AX activation after 48 hr. drug treatments**. U-2 OS cells were treated with 10 µM platinum compounds or 5 nM of ActD for 48 hours. The immunofluorescence intensity for γH2AX was then measured and percent positive nuclei were determined, where a positive threshold was determined by the 90th percentile of the untreated control. The data points represent the average percent positive nuclei across three separate biological replicates with standard deviation.

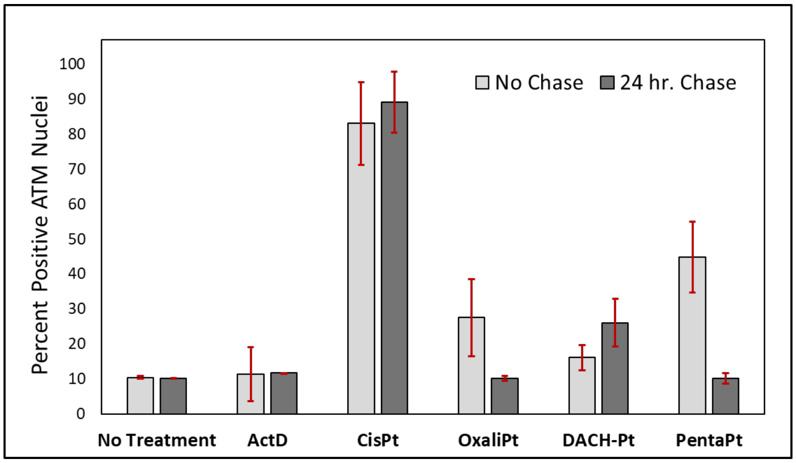


**Figure S2: Quantification of ATM activation after 5 hr. drug treatment and a 24 hr. drug free chase period.** A549 ells were treated with 10 µM platinum compounds or 5 nM of ActD for 5 hours. For the cells undergoing drug free chase, following treatment, drug-free media was replaced for 24 hours. The immunofluorescence intensity for pATM was then measured and quantified using the same method described in **Figure 4**. The data points represent the average percent positive nuclei with standard deviations across three separate biological replicates.

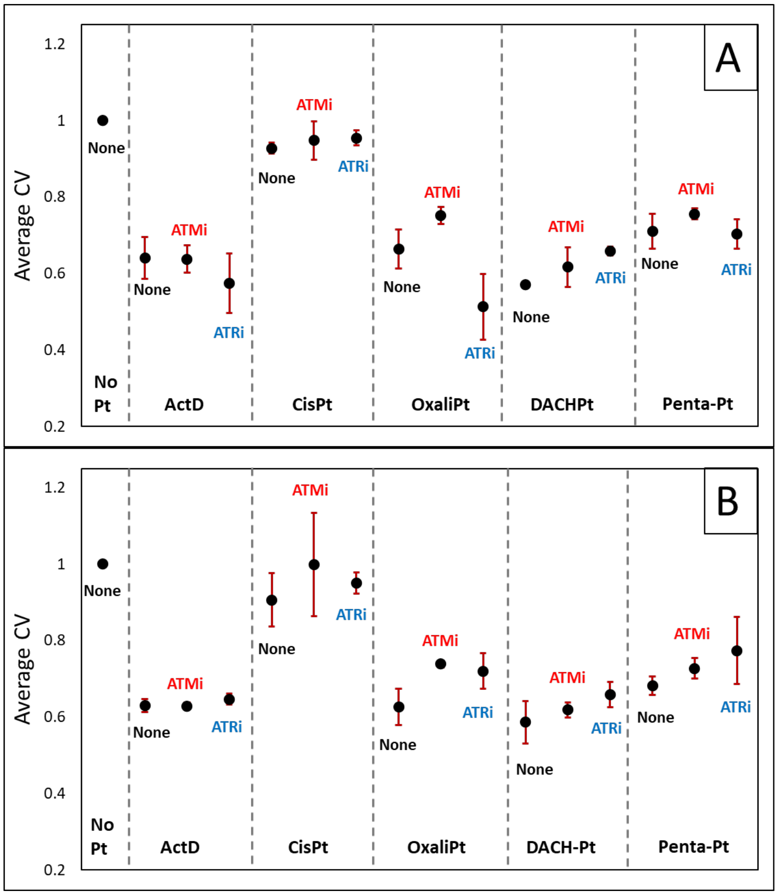


**Figure S3: NPM1 relocalization studies with ATM/ATR inhibitors at 5 hr. (A) and 3 hr. (B) drug treatment in A549 cells**. Cells were treated with 10 µM platinum compounds or 5 nM of ActD in the presence or absence ATMi (10 µM) and ATRi (2.5 µM). NPM1 immunofluorescence distribution was then quantified (Methods). Each point is the average CV value and standard deviation for 3 biological replicates

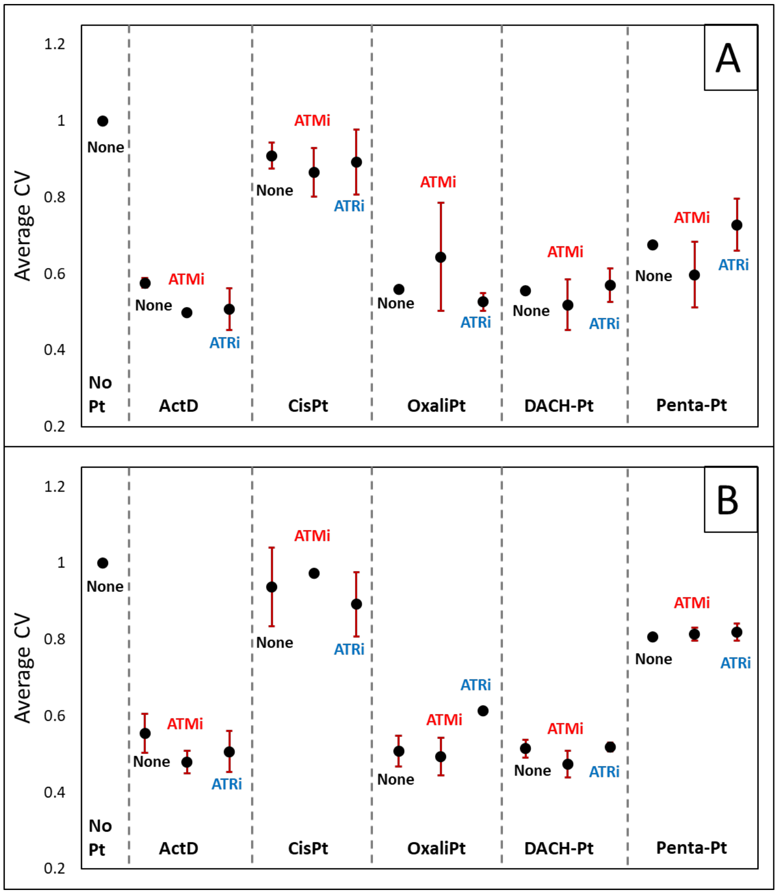


**Figure S4: NPM1 relocalization studies with ATM/ATR inhibitors at 5 hr. (A) and 3 hr. (B) drug treatment in U-2 OS cells.** Cells were treated with 10 µM platinum compounds or 5 nM of ActD in the presence or absence ATMi (10 µM) and ATRi (2.5 µM). NPM1 immunofluorescence distribution was then quantified (Methods). Each point is the average CV value and standard deviation for 3 biological replicates

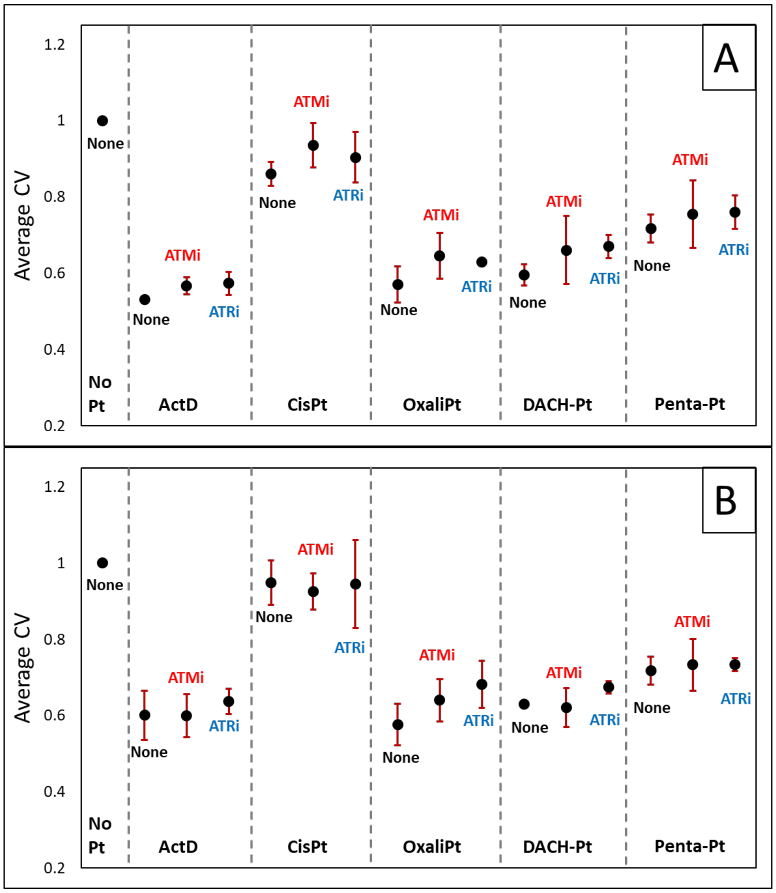


**Figure S5: NPM1 relocalization studies with ATM/ATR inhibitors at 5 hr. (A) and 3 hr. (B) drug treatment in HCT116 cells.** Cells were treated with 10 μM platinum compounds or 5 nM of ActD in the presence or absence ATMi (10 μM) and ATRi (2.5 μM). NPM1 immunofluorescence distribution was then quantified (Methods). Each point is the average CV value and standard deviation for 3 biological replicates

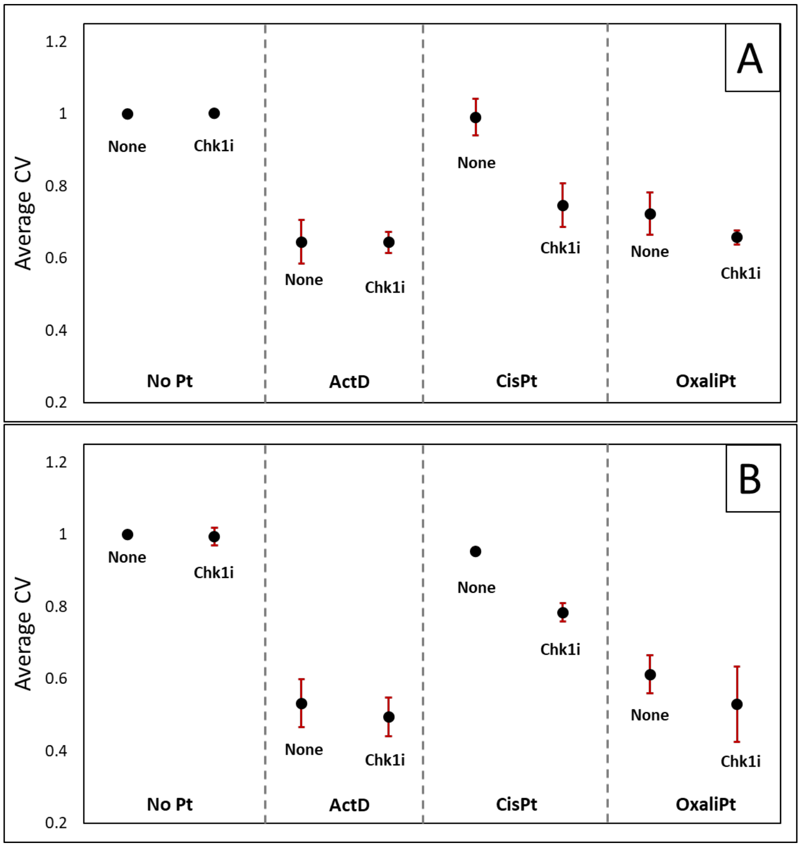


**Figure S6: NPM1 redistribution with Chk1 inhibition at 5 hr. (A) and 3 hr. (B) drug treatment in A549 cells.** Cells were treated with 10 µM platinum compounds or 5 nM of ActD for 24 hours in the presence or absence of Chk1i (1.0 mM). NPM1 immunofluorescence distribution was then quantified (Methods). Each point is the average CV value and standard deviation for 3 biological replicates.

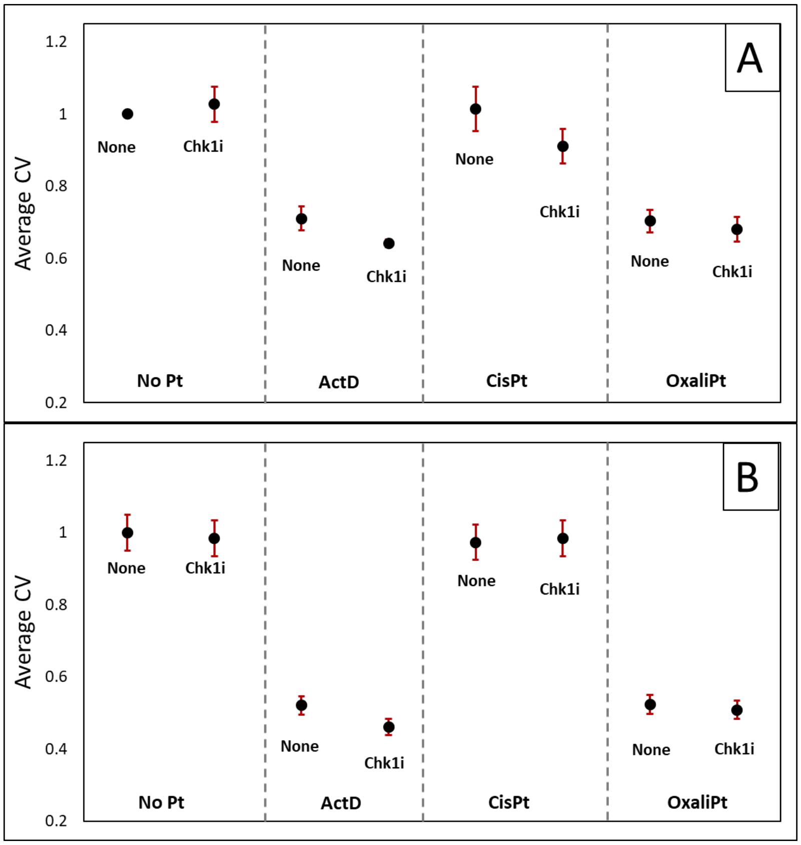


**Figure S7: NPM1 redistribution with Chk1 inhibition at 5 hr. (A) and 3 hr. (B) drug treatment in U-2 OS cells.** Cells were treated with 10 µM platinum compounds or 5 nM of ActD for 24 hours in the presence or absence of Chk1i (1.0 mM). NPM1 immunofluorescence distribution was then quantified (Methods). Each point is the average CV value and standard deviation for 3 biological replicates.

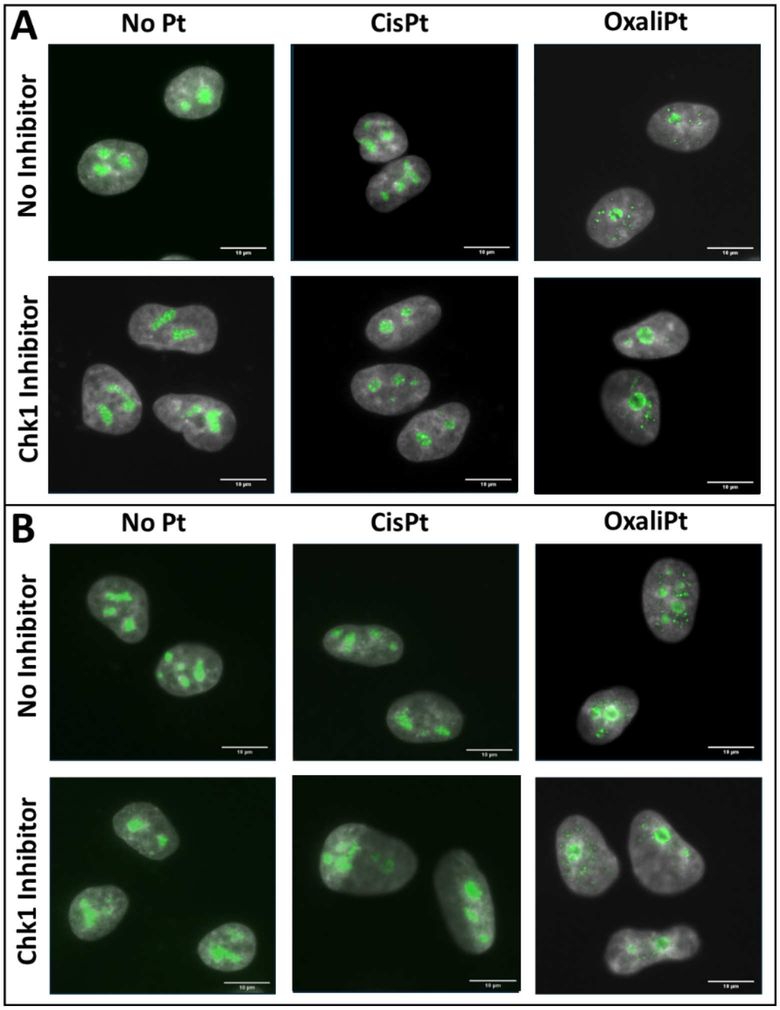


**Figure S8: Fibrillarin cap formation with Chk1i at 5 hr. drug treatment**. Cells were treated with 10 µM platinum compounds or 5 nM of ActD for 5 hours in the presence or absence of Chk1i (1.0 mM) in A549 (**A**) or U-2 OS (**B**) cell lines. Fibrillarin cap formation was then determine based on immunofluorescent images. Representative images for the no drug control and cells treated with **1** and **2** in the presence and absence of Chk1i are shown with fibrillarin (green) overlayed with DAPI (grey).

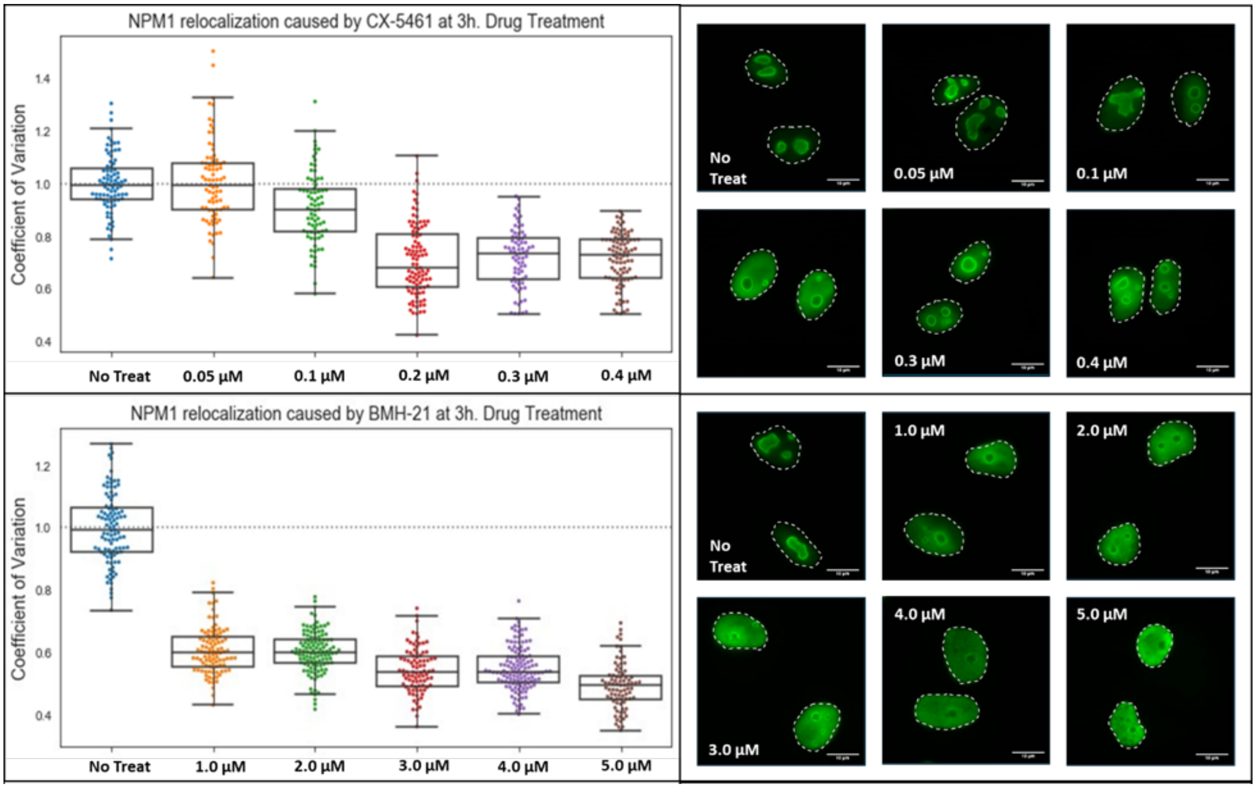


**Figure S9: NPM1 relocalization with small molecule nucleolar stress inducing compounds.** A549 cells were treated with various concentrations of CX-5461 (**A**) or BMH-21 (**B**) for 3 hours. NPM1 immunofluorescence distribution was then quantified (Methods). Representative cell images of NPM1 (green) are shown

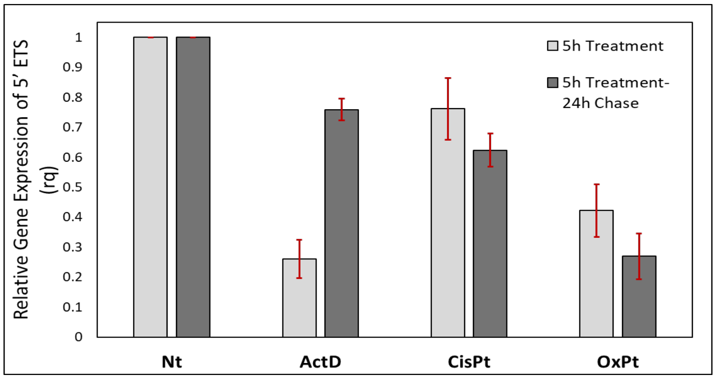


**Figure S10: rRNA transcription qPCR studies at 5hr. drug treatment and 24 hr. drug-free chase.** Cells were treated with 10 µM platinum compound or 5 nM ActD for 5 hrs. and then drug-free media was replaced for 24 hrs. The relative expression of 5' ETS was then determined (Methods) and plotted as an average with standard deviations from 3 biological replicates
